# Decoupled evolution of the *Sex Peptide* gene family and *Sex Peptide Receptor* in *Drosophilidae*

**DOI:** 10.1101/2023.06.29.547128

**Authors:** Ben R. Hopkins, Aidan Angus-Henry, Bernard Y. Kim, Jolie A. Carlisle, Ammon Thompson, Artyom Kopp

## Abstract

Across internally fertilising species, males transfer ejaculate proteins that trigger wide-ranging changes in female behaviour and physiology. Much theory has been developed to explore the drivers of ejaculate protein evolution. The accelerating availability of high-quality genomes now allows us to test how these proteins are evolving at fine taxonomic scales. Here, we use genomes from 264 species to chart the evolutionary history of Sex Peptide (SP), a potent regulator of female post-mating responses in *Drosophila melanogaster*. We infer that *SP* first evolved in the *Drosophilinae* subfamily and has followed markedly different evolutionary trajectories in different lineages. Outside of the *Sophophora-Lordiphosa*, *SP* exists largely as a single-copy gene with independent losses in several lineages. Within the *Sophophora-Lordiphosa,* the *SP* gene family has repeatedly and independently expanded. Up to seven copies, collectively displaying extensive sequence variation, are present in some species. Despite these changes, *SP* expression remains restricted to the male reproductive tract. Alongside, we document considerable interspecific variation in the presence and morphology of seminal microcarriers that, despite the critical role SP plays in microcarrier assembly in *D. melanogaster*, appear to be independent of changes in the presence/absence or sequence of SP. We end by providing evidence that SP’s evolution is decoupled from that of its receptor, SPR, in which we detect no evidence of correlated diversifying selection. Collectively, our work describes the divergent evolutionary trajectories that a novel gene has taken following its origin and finds a surprisingly weak coevolutionary signal between a supposedly sexually antagonistic protein and its receptor.

**Significance:** In insects, seminal fluid proteins (SFPs) induce dramatic changes in female behaviour and physiology. How this degree of male influence evolves remains a central question in sexual selection research. Here, we map the origin and diversification of the posterchild insect SFP, the *Drosophila* Sex Peptide (SP), across 264 Diptera species. We show that *SP* first evolved at the base of the subfamily *Drosophilinae* and followed markedly different evolutionary trajectories in different lineages, including accelerated change in sequence, copy number, and genomic position in the lineage leading to *D. melanogaster.* By contrast, we find only limited, uncorrelated change in the sequence of its receptor, SPR, arguing against a sexually antagonistic coevolutionary arms race between these loci on macroevolutionary time scales.

## Introduction

Female post-mating changes are a taxonomically widespread – if not general – phenomenon in internal fertilisers. Often mediated by non-sperm components of the male ejaculate, such as seminal fluid proteins, the female traits subject to post-mating plasticity are numerous and diverse. For example, immune systems can be modified (*Drosophila melanogaster*, 1 ; humans, 2), ovulation stimulated (camelids, 3), and dietary preferences shifted following copulation (crickets, 4 ; *D. melanogaster*, 5). Evolutionary biologists have a long-standing interest in post-mating changes as they bear intimate connections to reproductive success (e.g., 6–8), can form barriers to hybridisation (e.g., 9, 10), and, through the involvement of males in their induction, can act as a point of evolutionary tension between the fitness interests of males and females (e.g., 11–14). Indeed, post-mating changes have provided one of the centrepieces around which much of the discussion of interlocus sexual conflict has revolved, including broader consideration of the relative roles of conflict and sexual selection by female choice in shaping the evolution of reproductive traits (15–19).

Available data suggest that different taxa can use non-homologous proteins to induce common – or at least overlapping – phenotypic endpoints in mated females. To reduce female sexual receptivity, for example, the moth *Helicoverpa zea* uses pheromonostatic peptide (PSP, 20), the mosquito *Aedes aegypti* uses Head Protein 1 (HP-1, 21), and *D. melanogaster* uses Sex Peptide (SP, 22). Not only are these three proteins non-homologous to one another, but none have clear homologs in either of the other species’ genomes. This pattern suggests that regulators of female post-mating change might experience a high degree of evolutionary turnover, with new regulators evolving and old regulators being lost from populations through time. If so, the questions then are what evolutionary forces drive this turnover, how quickly does this process occur, and how are new regulators born? More fundamentally, it makes seminal proteins an exceptional model for studying how newly evolved, lineage-specific genes acquire and diversify their functions.

More than 60 years on from its discovery in chromatographic extracts of *D. melanogaster* accessory glands (23, 24), SP remains the best characterised insect seminal protein. Consisting of two exons separated by a 65bp intron, *D. melanogaster SP* (*DmelSP*) encodes a 36aa mature protein synthesised via a 55aa signal peptide-containing precursor (22, 25). DmelSP is produced in accessory gland main cells and secreted into the lumen, where it is stored on abundant, lipid-containing microcarriers – structures for which DmelSP acts as a key factor governing assembly and, once inside females, disassembly (26–28). Following transfer to females, DmelSP binds to sperm, a process mediated by a suite of additional seminal fluid proteins, and is transported into the female sperm storage organs (29–33). The gradual release of DmelSP from the surface of stored sperm continues to stimulate a wide range of post-mating changes, including shifts in memory formation and sleep patterns, elevating appetite and changing dietary preferences, reducing sexual receptivity, stimulating egg-laying, increasing aggression, and changing gut, metabolic, and immune activity (reviewed in 15). At least some of these changes, namely reduced sexual receptivity and stimulated egg-laying, are mediated by DmelSP binding to the Sex Peptide Receptor (SPR) in a subset of neurons that innervate the female reproductive tract (34–36). Different domains of mature SP appear to contribute selectively to different functions in *D. melanogaster*: the tryptophan-rich N-terminus binds to sperm and stimulates juvenile hormone synthesis (33, 37, 38), the hydroxyproline-rich mid-section elicits the innate immune response (39), and, through interactions with SPR, the disulphide bridge-containing C-terminus stimulates the core post-mating responses of increased oviposition and reduced sexual receptivity (40–42). Consequently, different portions of the *SP* coding sequence are likely to be evolving in response to different selective pressures.

*SP* is not the only member of its gene family present in *D. melanogaster.* This species also encodes the paralogous *Dup99b* with which SP shares a high degree of similarity in the amino acid sequence of the C-terminus (43). Both stimulate the core post-mating responses of increased oviposition and reduced sexual receptivity, but SP appears to be the ‘key player’ showing a higher binding affinity for the female reproductive tract and nervous system (41) and, in *in vitro* assays, activating SPR at lower concentrations than does Dup99b (36). There are further differences between the paralogs, too. While *SP* is expressed in accessory gland main cells, *Dup99b* is expressed in the ejaculatory duct (44). And, unlike SP, the N-terminus of Dup99b does not stimulate juvenile hormone synthesis (38). Thus, SP and Dup99b show partial redundancy but different sensitivities within one region of the protein and distinct activities in other regions, suggesting a degree of functional separation between the two paralogs.

SP occupies an important place in contemporary evolutionary biology, having emerged as one of the preeminent systems for experimental work on the genetic basis and fitness effects of sexual conflict (6, 45–47). However, comparative data on how SP sequence and function has evolved and diversified through time is sparse by comparison. Losses of *SP* have been reported in three *Drosophila* species (*D. grimshawi, D. albomicans,* and *D. mojavensis*) and the gene’s origin has been traced as far back as the most recent common ancestor of *D. virilis* and *D. melanogaster; SP* is apparently absent from mosquitoes and insect orders beyond Diptera (25, 48, 49). This contrasts with its receptor, *SPR*, which is deeply conserved among members of the Ecdysozoa and Lophotrochozoa, where it interacts with a similarly well conserved class of alternative ligands, the myoinhibitory peptides (MIPs)(48). MIP-SPR interactions are known to regulate diverse behaviours across species, including regulating larval settlement behaviour in marine annelids (50). In *Drosophila,* MIP-SPR interactions appear to be neither necessary nor sufficient for driving post-mating changes in females (48, 51), but they do fulfil other functions, including regulating sleep behaviour (52). Despite SP predating the group, several features of the SP-SPR system appear to be collectively restricted to the *melanogaster* species group, namely robust expression of *SPR* in the female reproductive tract, the ability of DmelSP to bind to female reproductive tract tissue, and a reduction in sexual receptivity upon injection of conspecific SP (25). Thus, despite the presence of SP orthologs beyond the group, many of the defining features of SP in *D. melanogaster* appear to be recently derived.

Taking advantage of newly available genomes for over 250 drosophilid species, here we report that *SP* is a drosophilid innovation that originated in the lineage leading to the *Drosophilinae* subfamily. We show that *SP* has subsequently followed markedly different evolutionary trajectories in different branches of the phylogeny, including lineage-accelerated evolution in sequence, copy number, and translocation frequency in the *melanogaster* group. Despite these changes, *SP* expression remains restricted to the male reproductive tract. We further fail to find support for the hypothesis that change in SP presence/absence or sequence is a significant driver of evolutionary change in microcarrier morphology. Finally, we fail to find a signal of coevolution between *SP* and the receptor through which it induces many of its effects in females, *SPR,* arguing against a sexually antagonistic coevolutionary arms race between these loci on macroevolutionary time scales.

## Results

### *Sex Peptide* first evolved in the *Drosophilinae* subfamily

To pinpoint the origin of *SP,* we designed a pipeline to identify *SP* orthologs in whole-genome sequences. Our approach combined reciprocal blast of the *D. melanogaster* SP C-terminus sequence with protein sequence, gene structure, and synteny analysis. As in a previous study (48), we failed to detect *SP* in a genome from the mosquito *Aedes aegypti,* and additionally failed to detect *SP* in genomes from two calyptrate members of the Brachycera suborder to which *Drosophila* belongs: *Musca domestica* (53) and *Glossina morsitans* (54). Within the Acalyptratae, we found that *SP* was restricted to the *Drosophilidae –* specifically to the subfamily *Drosophilinae* Rondani (Figure 1). Within the *Drosophilinae*, *SP* was present in genomes from several species that predate the *Drosophilini,* the tribe that includes all members of the *Drosophila* genus (55). These non-*Drosophilini* species included members of the *Colocasiomyini* tribe, including species within the *Scaptodrosophila, Lissocephala, Chymomyza,* and *Colocasiomyia* genera, but several secondary losses of SP were apparent. We failed to detect *SP* in genomes from 10 members of the other *Drosophilidae* subfamily, the *Steganinae* Hendel, or members of several closely related outgroups, namely the genera *Liriomyza, Cirrula, Ephydra,* and *Diastata.* We therefore infer that *SP* first evolved in the *Drosophilinae* subfamily.

**Figure 1.**
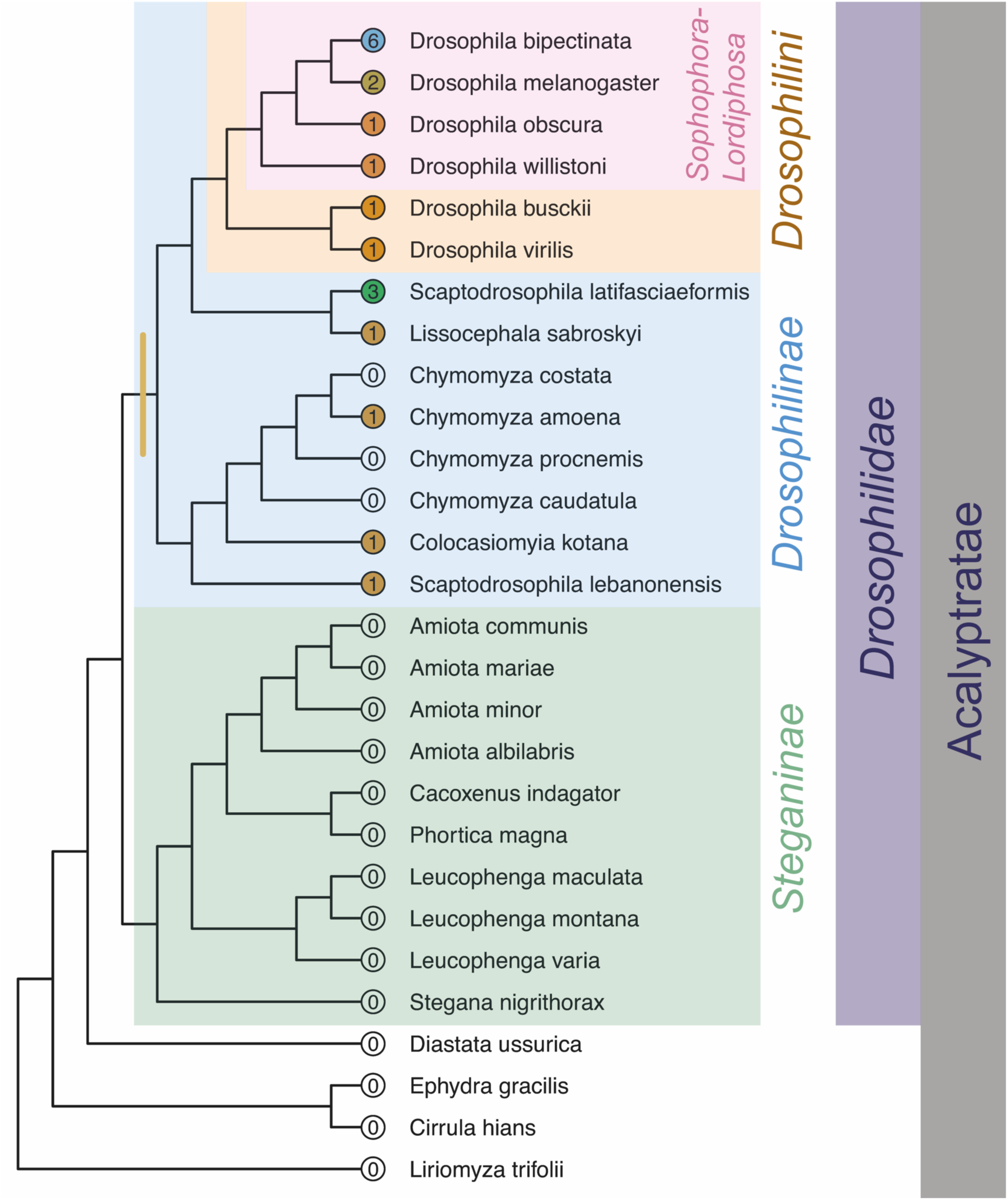
*SP* first evolved in the *Drosophilinae* subfamily. A phylogeny of species from across the *Drosophilidae* and closely related lineages. The number of *SP* genes detected is given in coloured circles at the tip of each branch. A brown bar denotes the branch on which we infer *SP* to have first evolved.

### *Sex Peptide* has been repeatedly lost and rarely duplicated outside of the *Sophophora-Lordiphosa* radiation

Within our phylogenetic sample, the *Drosophilini* splits into two lineages. The first contains the *Lordiphosa* genus and *Sophophora* subgenus (which we collectively refer to as the ‘*Sophophora-Lordiphosa* radiation’), the latter of which includes the *melanogaster, obscura, willistoni,* and *saltans* groups (see *SI Appendix Fig. S1* for an overview of the *Sophophora* taxonomic terminologies used in this paper). The second lineage includes, among others, the genera *Scaptomyza* and *Zaprionus* and the Hawaiian*, virilis, repleta, immigrans, cardini,* and *quinaria* groups (the *Drosophila* genus is paraphyletic (55, 56)).

Outside of the *Sophophora-Lordiphosa* radiation, we observed several features of *SP’*s evolution. First, *SP* has been repeatedly and independently lost—four times in our phylogenetic sample. Once in a monophyletic lineage of 29 species covering the *annumilana, bromeliae*, *nannoptera*, *mesophragmatica*, and *repleta* groups (Figure 2; *SI Appendix, Fig. S2*). The other three separately covered all 55 Hawaiian species in our dataset (Figure 2; *SI Appendix, Fig. S3*), a monophyletic lineage within the *Scaptomyza* (Figure 2; *SI Appendix, Fig. S3*), and a species-specific loss in *Hirtodrosophila duncani* (Figure 3). The second trend was that duplications of *SP* were rare. Among the 53 non-*Sophophora-Lordiphosa* species in the *Drosophilini* in which we did detect *SP,* all but one had just a single copy (Figure 2). The exception was *D. paramelanica* (*melanica* group) in which we detected a tandem duplication, with the two copies sharing 100% identity in predicted protein sequence. Outside of the *Drosophilini,* only *Scaptodrosophila latifasciaeformis* showed an expansion in *SP* copy number, bearing 3 tandemly arranged copies that diverged from one another in predicted protein sequence. The third trend was that *SP* rarely translocated to new genomic locations. Where *SP* was detected in non-*Sophophora-Lordiphosa* species in the *Drosophilini*, in all but two we found that it mapped to a syntenic neighbourhood on Muller element D, which we call ‘Muller D1’, that contained orthologs of *FoxK, NaPi-III,* and *mRpL2.* The exceptions were *Hirtodrosophila trivittata,* where *SP* mapped to a distinct neighbourhood on Muller element D that contained orthologs of *bruno3, CG3349,* and *CG17173,* and *D. repletoides*, in which *SP* mapped to a neighbourhood on Muller element B containing orthologs of *halo* and *haf*. Outside of the *Drosophilini,* we also detected *SP* in the Muller D1 neighbourhood in *S. lebanonensis* and *S. latifasciaeformis,* suggesting that this may be the ancestral position of *SP*.

**Figure 2.**
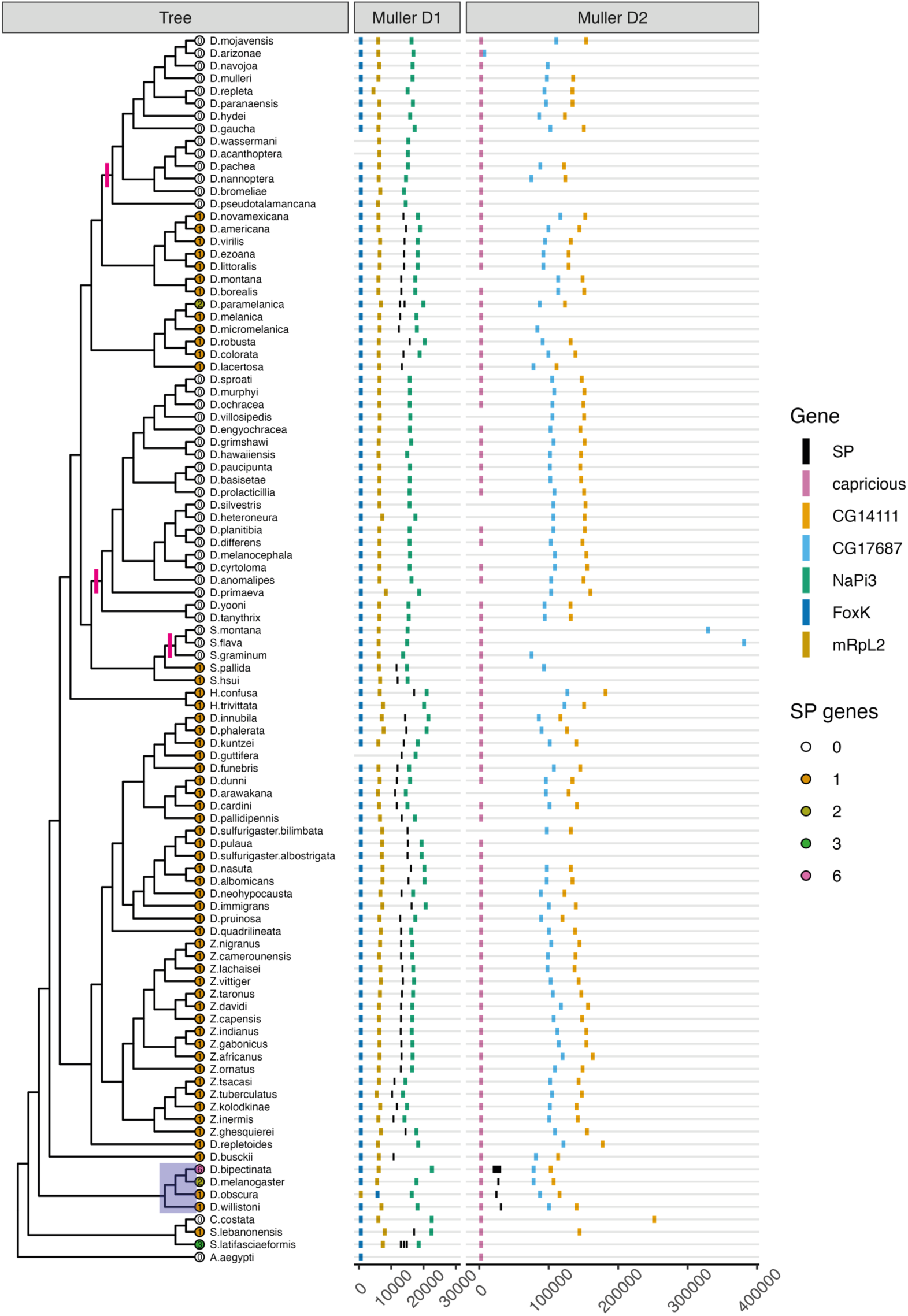
*Sex Peptide* family genes predate the *Drosophilini* and have been repeatedly lost outside of the *Sophophora-Lordiphosa* radiation. This figure focuses on the non-*Sophophora-Lordiphosa* members of the *Drosophilini* (see Figure 1 and *SI Appendix, Fig. S1* for overviews of drosophilid taxonomy). A selection of *Sophophora* species, shaded in blue, are included for comparison. Also included are four non-*Drosophilini* dipterans: *Aedes aegypti* and three non-*Drosophilini* members of the *Drosophilinae* subfamily: *Chymomyza costata, Scaptodrosophila lebanonensis,* and *S. latifasciaeformis.* The number of *SP* genes detected in a representative of each species’ genome is given at the tree tips. Losses are marked with a pink bar. For each species, the structures of two syntenic gene neighbourhoods are plotted. The first, Muller D1, is the canonical neighbourhood in which *SP* genes are detected outside of the *Sophophora-Lordiphosa.* The second, Muller D2, is the canonical position in the *Sophophora-Lordiphosa.* Positions of each gene are given relative to the first gene in the neighbourhood (*FoxK* or *capricious*). Absence of a flanking neighbourhood gene (*e.g., FoxK* in *D. wassermani*) doesn’t necessarily mean the gene has been lost – it more likely means that a contig breakpoint fell within the neighbourhood. Note that *SP* in *Hirtodrosophila trivittata* and, independently, *D. repletoides,* has translocated out of the Muller D1 neighbourhood. See *SI Appendix, Figs S2 and 3* for expanded views of the losses in the *repleta* and *Hawaiian* groups, respectively.

**Figure 3.**
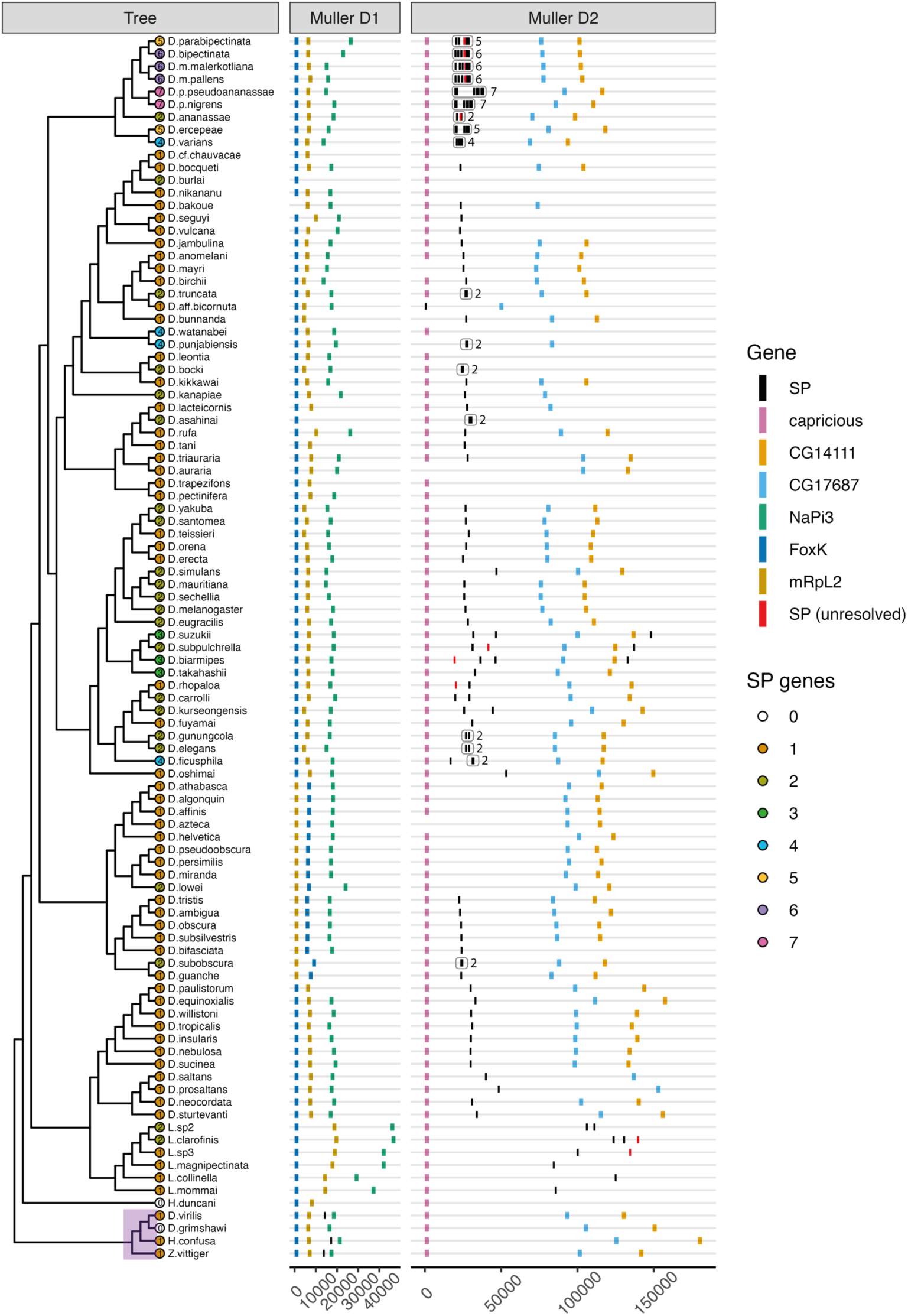
*Sex Peptide* copy number is markedly more variable in the *Sophophora-Lordiphosa* radiation than in other branches of the phylogeny. This figure focuses on the *Sophophora-Lordiphosa* radiation to which *D. melanogaster* belongs. Four non-*Sophophora-Lordiphosa* drosophilids, shaded in purple, are included as outgroups. The structures of the Muller D1 and Muller D2 neighbourhoods are plotted as in Figure 2. Missing flanking genes are likely indicative of contig breakpoints falling within the neighbourhood. The exceptions are the *Lordiphosa* species, where substantially elevated intergenic distances meant that the whole neighbourhood would not fit within the plot limits. Unresolved SP genes, shown in red, indicate genes that passed the reciprocal blast criteria and fell within one of the conserved *SP*-containing gene neighbourhoods but where a SP-like amino acid sequence couldn’t be resolved (*e.g.,* due to a premature stop codon, as in the case of *D. rhopaloa*). Note that all members of the *obscura* group have an inversion that flips the relative positions of *FoxK* and *mRpL2* in the Muller D1 neighbourhood. In a number of cases, some or all copies of *SP* were found to have translocated outside of the Muller D1 and Muller D2 neighbourhoods (an *obscura* group lineage, the *melanogaster* subgroup, *D. kanapiae, D. takahashii*, and *D.eugracilis*; summarised in *SI Appendix Fig. S5*). In the shorter read *montium* subgroup assemblies, short contigs meant that in some species we couldn’t identify the neighbourhood in which *SP* was located. This was the case for some *SP* genes in *D. cf. chauvacae, D. burlai, D. leontia, D. nikananu, D. pectinifera, D. punjabiensis,* and *D.watanabe*i. The *SP* genes in *D. auraria* and *D. trapezifons* could be mapped to the Muller D2 neighbourhood based on flanking sequence around the *SP* gene, but the *SP*-containing contigs were too small to include any of the neighbourhood genes.

### *Sex Peptide* has repeatedly duplicated in the *Sophophora-Lordiphosa* radiation

Within the *Sophophora-Lordiphosa, SP* has followed a markedly different evolutionary trajectory. For one, and despite denser taxon sampling in this part of the phylogeny, we detected at least one copy in all species sampled—no species was entirely without *SP.* Second, we detected a clear uptick in the frequency of duplication (Figure 3). In the earlier branching lineages, we detected apparently independent duplications within a sublineage of the *Lordiphosa,* in *D. subobscura* (obscura group; see also 57), and in *D. lowei* (*obscura* group). Within the *melanogaster* group we found much greater variability in *SP* copy number, consistent with repeated, lineage-specific expansions and contractions of gene family size. This variation was greatest in the “Oriental” lineage, which includes *D. melanogaster,* and *ananassae* subgroup. As many as 7 tandemly arranged copies were present in some *ananassae* subgroup species (Figure 4).

**Figure 4.**
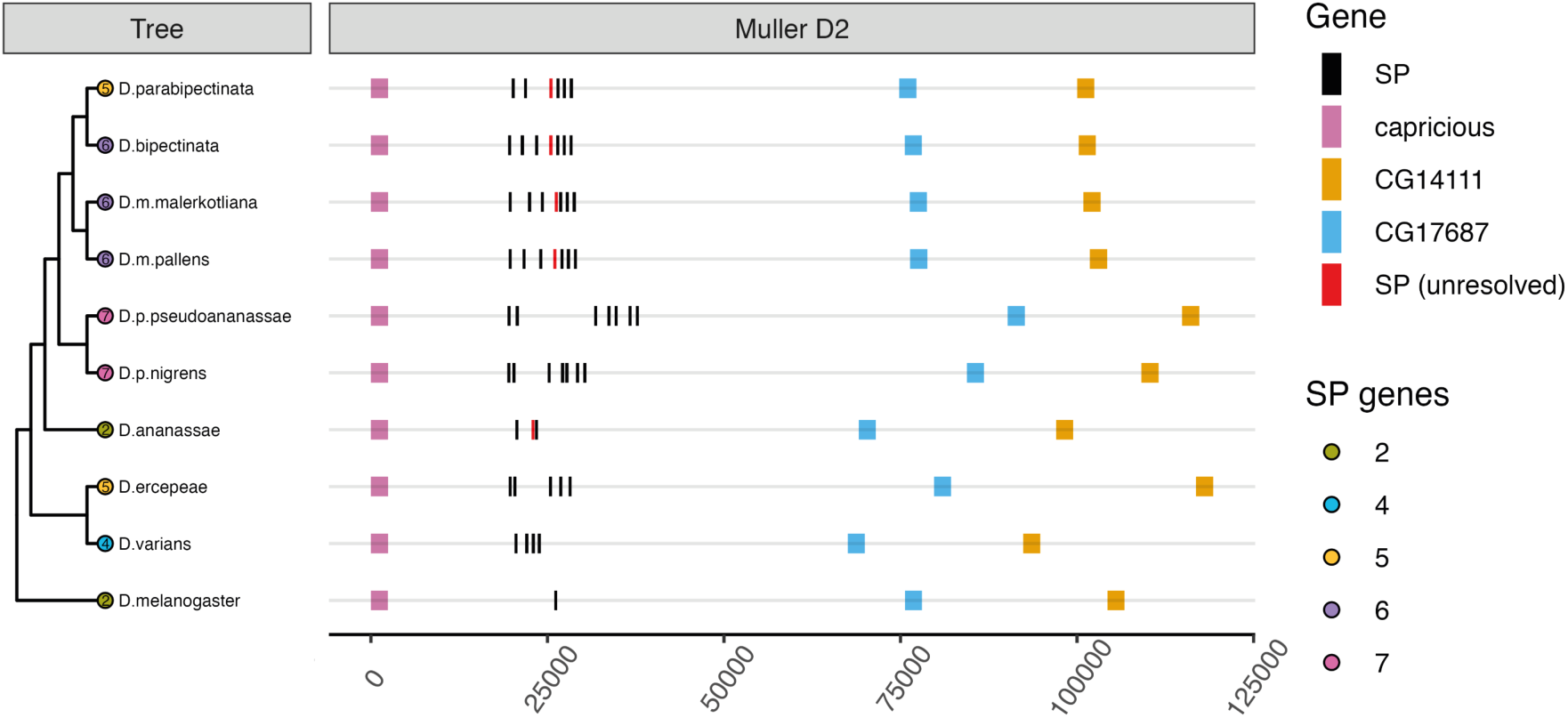
Repeated duplication of *Sex Peptide* genes in the *ananassae* subgroup. A phylogeny of the *ananassae* subgroup species used in this study, with *D. melanogaster* as an outgroup. The number of *SP* genes identified in each species is given at the tip of each branch. ‘Unresolved’ SP sequences, shown in red, are those which passed the reciprocal blast tests and fell within the syntenic Muller D2 gene neighbourhood but for which we could not resolve an SP-like protein sequence (*e.g.,* due to a premature stop codon). The structure of the neighbourhood is plotted on the right-hand side of the figure. Note that one of *D. melanogaster’s SP* copies, *Dup99b,* falls outside of the Muller D2 neighbourhood.

To resolve the evolutionary relationships between *SP* paralogs within the *Sophophora-Lordiphosa* radiation, we constructed a tree of the predicted SP protein sequences (*SI Appendix, Fig. S4*). The tree supports numerous recent duplications affecting single species or species pairs, including in *D. elegans/D.gunungcola, D. takahashii, D.ficusphila, D. punjabiensis/D. watanabei, D. kanapiae,* and *D. subobscura*. The tree also suggests that there have been three separate expansions of *SP* copy number within the *ananassae* subgroup, one in the *bipectinata* complex and another in each of *D. varians* and *D. ercepeae* (*SI Appendix, Fig. S4*, coloured orange and blue, respectively), likely from an ancestral starting point of the two copies seen in *D. ananassae.* We note, however, that the sequence similarity that we observe between putative paralog pairs may be driven instead by concerted evolution.

### Frequent translocations of *Sex Peptide* genes within the *Sophophora* subgenus

At the base of the *Sophophora-Lordiphosa* radiation, *SP* appears to have translocated from the Muller D1 neighbourhood to a new neighbourhood ∼2.1Mb away on the same Muller element (Figure 3). This syntenic neighbourhood, which we refer to as ‘Muller D2’, contains orthologs of *capricious, CG14111,* and *CG17687.* Despite translocation, the configuration of the ancestral Muller D1 gene neighbourhood remains intact in the *Sophophora-Lordiphosa*. Thus, the mechanism of translocation did not lead to the breakup of the Muller D1 neighbourhood via a larger scale rearrangement. Several further translocations are then present (summarised in *SI Appendix, Fig. S5*; *e.g.,* in the *obscura* group, *SI Appendix, Fig. S6*). Each of *D. suzukii, D. subpulchrella,* and *D. biarmipes*, which form a monophyletic clade within the “Oriental” lineage, bear SP copies in the canonical Muller D2 position with an additional copy just the other side of *CG14111* within the same neighbourhood (Figure 3). The protein tree supports a *Dup99b* identity for these copies that have skipped to the other side of *CG14111* (*SI Appendix, Fig. S4*, shown in pink). In *D. takahashii,* the sister species to this clade, two of the three SP genes we detected mapped to a neighbourhood on *D. melanogaster* Muller element B that contained *NLaz, robo2,* and *CG14346* orthologs. The protein tree also supports a *Dup99b* identity for these translocated copies (*SI Appendix, Fig. S4*, shown in pink). However, given the low support for internal nodes, we were not able to accurately determine the timing of the initial duplication that gave rise to separate *SP* and *Dup99b* copies.

In *D. melanogaster, SP* falls within the Muller D2 neighbourhood while its paralog *Dup99b* maps to Muller element E in a neighbourhood that contains *dmrt99b, gycalpha99b,* and *CG34296.* This arrangement appears to be ancestral to the *melanogaster* subgroup (*SI Appendix, Fig. S7A*,*B*). The losses of an *SP* gene within a subset of species in this subgroup, namely *D. teissieri* (strain CT02; present in 273.3)*, D. orena,* and *D. erecta,* affect the Muller element E *Dup99b* copy, rather than the Muller D2 *SP* copy. In *D. eugracilis,* the *melanogaster* subgroup’s sister species, *SP* is present in the Muller D2 neighbourhood, with a second copy in a different position on Muller element E that contains orthologs of *SmD2, CG18048,* and *Hr83*. Despite the lack of synteny, the protein tree supports a *Dup99b* identity for this translocated copy (*SI Appendix, Fig. S4*).

### Male reproductive tract-biased expression is a conserved feature of *SP* genes

Our identification of *SP* genes was based on gene sequence data and synteny, leaving open the question of whether and where they are expressed. For 19 species, we were able to test for expression using RNA-seq datasets available through NCBI. 38 of the 42 *SP* genes we detected across the 19 species were expressed, although many were un- or incorrectly annotated (*e.g.,* as long non-coding RNAs) in the reference genomes (see *SI Appendix*). All 4 of those that weren’t expressed lacked SP-like protein sequences due to point mutations affecting either the start codon or introducing premature stop codons, suggestive of pseudogenisation (*e.g.,* in *D. rhopaloa*; *SI Appendix, Fig. S8*). Of the 38 *SP* genes we found to be expressed, all showed strongly male-biased expression, including in the early branching *D. busckii* (*SI Appendix, Fig. S9*). The one exception was detection of appreciable *SP* expression in a single *D. simulans* female sample, which was due to sample contamination or mislabelling (*SI Appendix, Fig. S10*).

In all 10 species where we had tissue-specific expression data, including the distantly related *D. virilis,* we observed clearly enriched expression of *SP* family genes in the male reproductive tract (*SI Appendix, Fig. S11*). Where datasets were available for sub-portions of the male reproductive tract, expression was generally substantially higher in samples labelled as accessory gland or non-gonadal reproductive tissues than in samples labelled as testes. The extent of testes expression was variable between samples and between species, perhaps reflecting varying degrees of contamination between these closely associated tissues during dissection. Based on these data, we conclude that male reproductive tract-biased expression is a conserved feature of the *SP* gene family and is therefore likely the ancestral expression pattern within the *Drosophilini*.

### Accelerated evolution of Sex Peptide proteins in the *Sophophora-Lordiphosa* radiation

Aligning 233 SP sequences from 148 genomes for the species shown in Figure 2 and 3, we find that the C-terminus, which is responsible for stimulating post-mating responses in *D. melanogaster* (33, 42), is highly conserved both inside and outside of the *Sophophora-Lordiphosa* radiation (*SI Appendix, Fig. S12A-C*). Several residues in this region, including the disulphide bond forming cysteine residues, are present in almost all SP sequences in our dataset: within the consensus sequence KWCRLNLGPAWGGRGKC, W2, C3, G8, P9, G12, G13, and C17 are each conserved in >97% of sequences (*SI Appendix, Fig. S12A*). In contrast, the mid-section, which has been implicated in stimulating innate immune responses (39), and the N-terminus (following cleavage of the signal peptide), which is responsible for binding to sperm and stimulating juvenile hormone synthesis (33), showed quite limited sequence conservation, suggesting more rapid evolutionary change.

The predicted length of SP proteins showed elevated variability in the *Sophophora-Lordiphosa* radiation (*SI Appendix, Fig. S12D*). These differences held after *in silico* cleavage of predicted signal peptides and were largely due to the introduction of additional amino acids upstream of the post-mating-response-stimulating C-terminus. For 38 genes across 19 species, we were able to use the RNA-seq data to validate our annotation of exon/intron boundaries. In all 38 cases our predicted boundaries matched those derived from the RNA-seq data. The expression data alone therefore supports a change in pre-cleavage SP sequence length between *e.g., D. virilis* (47aa) and *D. rhopaloa* (60aa), as well as *D. bipectinata* expressing a set of 6 SP proteins of variable length (46aa, 49aa, 54aa, 62aa, 68aa, 72aa). Moving on to SP protein sequence, a PCA generated from substitution matrix scores showed a high degree of dispersion among *Sophophora-Lordiphosa* orthologs relative to those from outside of the radiation (*SI Appendix, Fig. S12E-G*). This included clear separation from the remaining sequences of SP – but not Dup99b – orthologs from the “Oriental” lineage (except for the most basal species we sampled from this lineage, *D. oshimai*). Their distinct clustering may be driven by their N-terminus and midsection sequences, which showed limited conservation with those of other SP proteins. More generally, the high degree of dispersion between sequences in the *Sophophora-Lordiphosa,* and particularly the *ananassae* subgroup, points to a high degree of sequence diversity within this lineage.

### Microcarrier morphology is not clearly linked to the copy number of *Sex Peptide*

We next wanted to explore functional consequences of the diversity in the phylogenetic distribution and sequence of *SP* genes. Recently, SP was shown in *D. melanogaster* to be a key factor influencing the assembly, disassembly, and morphology of microcarriers, lipid-based structures that appear to store and traffic seminal fluid proteins (28). Because of this relationship between SP and microcarrier structure, it was suggested that variation in SP sequence might be associated with inter-specific variation in microcarrier morphology (28).

Using the neutral lipid-specific dye LipidTox, which has previously been used to stain microcarriers (28), we sought to examine the relationship between SP and microcarrier structure on two levels. The first was to ask whether variation in *SP* copy number is associated with a shift in microcarrier morphology. For this, the *ananassae* subgroup provides an ideal system, given that its constituent species encode between 2 and 7 *SP* copies. Within the *bipectinata* species complex, all of the species that we looked at (*D. parabipectinata, D. bipectinata, D. m. malerkotliana, D. m. pallens, D. p. nigrens*), which each encode between 5 and 7 *SP* copies, showed small, globular microcarriers (Figure 5A-F), similar to those seen in the *obscura* group (28). Curiously, however, and unlike those seen in the *obscura* group, these microcarriers appeared to carry a central indentation reminiscent of the biconcave disk shape of human red blood cells. This morphology was clearly distinct from that of *D. melanogaster* microcarriers, which appear as a heterogeneous mix of fusiform, ellipsoid, and thread-like structures (Figure 5L). The *bipectinata* complex morphology was also distinct from those of three other *ananassae* subgroup species: *D. ananassae* (2 copies), which had thread-like and spiral or doughnut shaped microcarriers (Figure 5G); *D. ercepeae* (5 copies), which had thread-like microcarriers (Figure 5H); and *D. varians* (4 copies), which displayed a highly divergent organization of the lumen’s contents (Figure 5I,J). In *D. varians*, LipidTox appeared to be excluded from vacuolar structures that were filled with small, weakly stained droplets. The vacuoles appeared larger in the proximal region of the gland, suggesting that they may fuse as they move towards the ejaculatory duct (Figure 5J). We observed a similar pattern in the three-copy-encoding *D. takahashii*, a non-*ananassae* subgroup species, although here the LipidTox staining was negligible (Figure 5N). This contrasted with *D. takahashii*’s close relative *D. biarmipes,* which despite also encoding three copies showed a unique staining pattern of strongly stained, tiny microcarriers that appeared to aggregate (Figure 5M). The microcarriers of *D. biarmipes* adopted a conformation reminiscent of *D. melanogaster’s* following transfer to the female reproductive tract as they begin to break down into smaller puncta (28). The conformation observed in *D. biarmipes* appears to be a derived state as the more distantly related *D. carrolli* (Figure 5O) bears microcarriers that more closely resemble those of *D. melanogaster, D. sechellia,* and *D. simulans* (Figure 5L; 23). Consequently, while the *melanogaster* group shows remarkable diversity in both microcarrier morphology and *SP* copy number, there appears to be no clear relationship between them.

**Figure 5.**
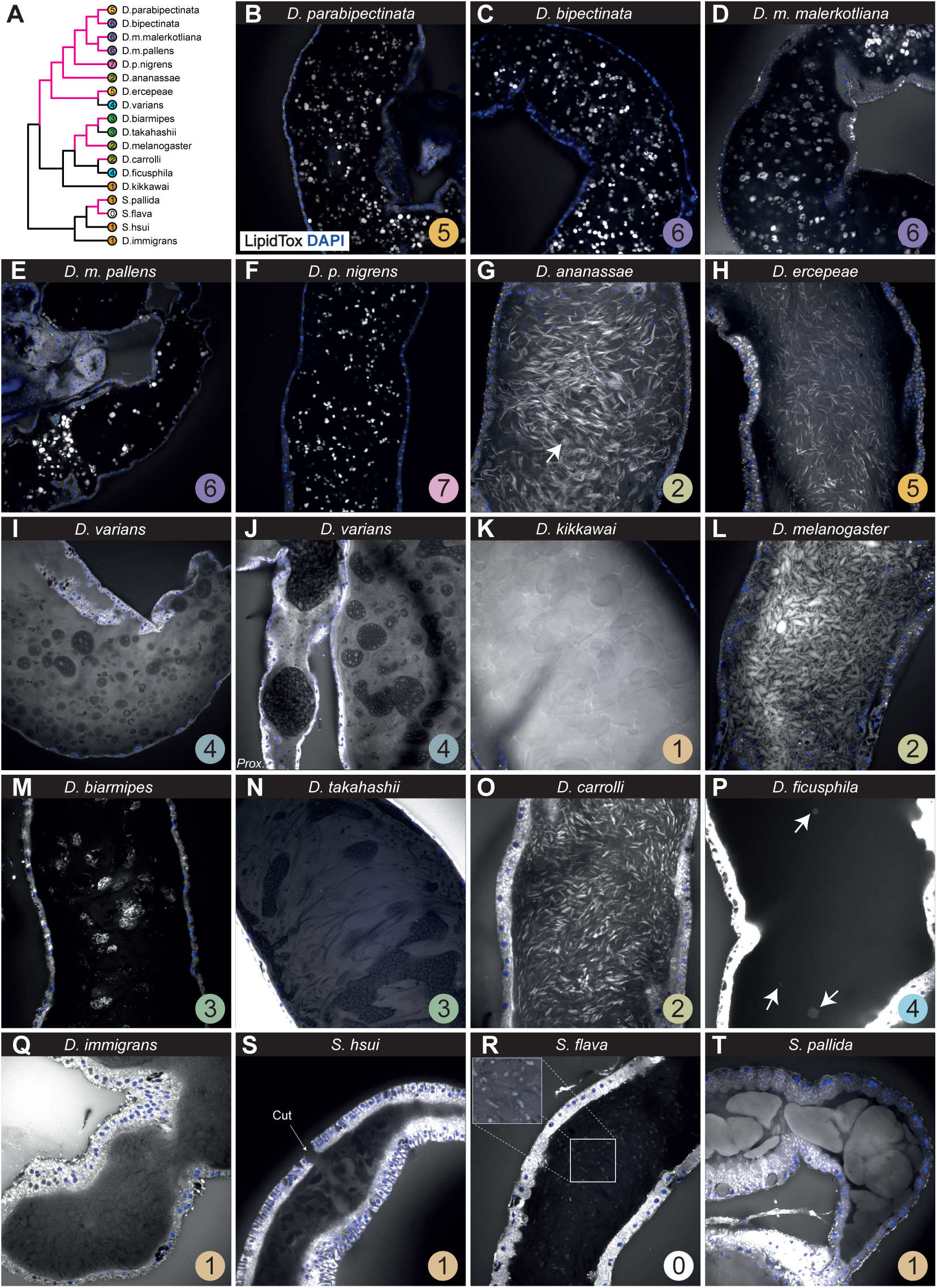
*Sex Peptide* is neither necessary nor sufficient for microcarriers. (A) A phylogeny of all species included in this figure. Branches coloured pink indicate species shown in B-T that demonstrate canonical (*i.e., D. melanogaster*-like) staining with LipidTox, a neutral lipid-specific dye used to selectively stain microcarriers (28). (B-T) Accessory glands stained with LipidTox and the nuclear stain DAPI (blue). The circled number in the bottom right-hand corner of each panel indicates the number of *Sex Peptide* copies we detect in each species. (G) The arrow is highlighting a spiral/doughnut shaped microcarrier, a shape which is rare in comparison to the more common thread-like conformation in this species. (J) Prox. refers to the proximal region of the gland, *i.e.* the region that connects to the ejaculatory duct. (P) Arrows point to the ambiguous, sparse, and weakly stained material we observed in *D. ficusphila* glands. (S) An arrow points to a cut in the glandular epithelium, which was made to enhance dye penetration. In each case, glands were co-stained in the same well with those from *D. melanogaster* to act as a positive control.

### Detection of microcarrier-like, but LipidTox^-^, structures within and beyond *Sophophora*

Previous staining of accessory glands from the single-copy-encoding *D. virilis* demonstrated that a copy of *SP* is not sufficient for LipidTox-stained microcarriers (28)*. D. virilis* instead displayed a more uniform ‘flocculence’ within the gland’s lumen that showed little evidence of LipidTox staining. We observed a similar flocculent arrangement in the single-copy-encoding *D. immigrans* (Figure 5Q). We also observed an essentially microcarrier-free glandular lumen in the four-copy-encoding *D. ficusphila* (Figure 5P). In this species we observed only a handful of weakly stained structures per gland, the rarity and structural inconsistency of which renders their classification as microcarriers doubtful. Alongside these cases, we detected instances of microcarrier-like, ellipsoid structures that failed to take up LipidTox in several species from diverse parts of the drosophilid tree, namely the *montium* subgroup species *D. kikkawai* (Figure 5K) and the non-*Sophophora* species *Scaptomyza hsui* (Figure 5S). All four of these species – *D.immigrans, D. ficusphila, D.kikkawai,* and *S. hsui –* each encode at least one *SP* copy, providing further support for the claim that a copy of *SP* is not sufficient for LipidTox^+^ microcarriers.

### Microcarriers predate the *Sophophora,* but copies of *Sex Peptide* are neither necessary nor sufficient for their presence

Staining glands from a species that we identified as having lost the *SP* gene*, S. flava,* we observed small, globular microcarriers reminiscent of those from the *obscura* group, albeit weaker in their staining (Figure 5R). Thus, *SP* is not necessary for microcarriers. Moreover, the previous complement of species that had been stained suggested that LipidTox^+^ microcarriers were confined to the *obscura* and *melanogaster* groups. We now show that they are present outside the *Sophophora*.

To better understand the distribution of microcarriers within the *Scaptomyza,* we also looked at *S. flava*’s single-copy-encoding sister species, *S. pallida*. This species showed strong LipidTox staining, but the pattern was unlike any other species we looked at (Figure 5T). Rather than the lumen being filled with large numbers of small microcarriers with well-defined shapes, the *S. pallida* lumen was filled with substantial clouds of stained secretion that in many cases spanned the full diameter of the gland’s internal space. This pattern was reminiscent of that observed in repeatedly mated *SP* null – but not wild-type – *D. melanogaster* males (28). Thus, in the presence of an *SP* ortholog we observe in *S. pallida* an apparent phenocopying of an *SP* null conformation, further evidence that microcarrier morphology may, at a broad taxonomic scale, be largely decoupled from evolutionary change in *SP*.

### No clear signal of episodic diversifying selection in SPR sequence among drosophilids

If SP is coevolving with the G-protein coupled receptor (GPCR) through which it induces many of its effects, SPR (36), then we might predict that bursts of evolutionary change in *SP* copy number and protein sequence correlate with similar bursts of change in SPR. Resolving *SPR* sequences from 193 genomes (the species shown in Figures 2 and 3), we failed to detect a single instance of duplication, suggesting that the mode of evolutionary change is decoupled between *SP* and *SPR.* For 5 species, including two that had lost *SP* (*S. flava* and *S. montana*), we failed to resolve an SPR sequence (*SI Appendix, Fig. S13*). Thus, while there is no phylogenetically repeatable association in the copy number of the two genes, the loss of one can be accompanied by the loss of the other.

At the sequence level, we found that evolutionary change in SPR is concentrated in the extracellular N-terminus domain, perhaps consistent with evolution under relaxed selection (see *SI Appendix, Supp. Text*), with the remainder of the protein sequence showing much stronger conservation (Figure 6A-C; *SI Appendix, Fig. S13*). PCA suggested that while the degree of diversity among SPR sequences was apparently higher in the *ananassae* subgroup and “Oriental” lineages relative to the *montium* subgroup, the overall diversity did not appear markedly elevated in these lineages compared to the full spectrum of SPR sequences (Figure 6D, E). After removing the poorly conserved N-terminus region, we found no evidence for episodic diversifying selection in *SPR* sequences using branch- (aBSREL, 58) or gene- (BUSTED, 59) level tests for selection. This held whether we formally tested all branches, only those in the *Sophophora-Lordiphosa* radiation, or only those outside of the *Sophophora-Lordiphosa* radiation (see *SI Appendix* for how to view the full output of these analyses). Thus, we fail to find evidence of a burst in evolutionary change in *SPR* that correlates with the lineage-accelerated evolutionary changes we record for *SP*.

**Figure 6.**
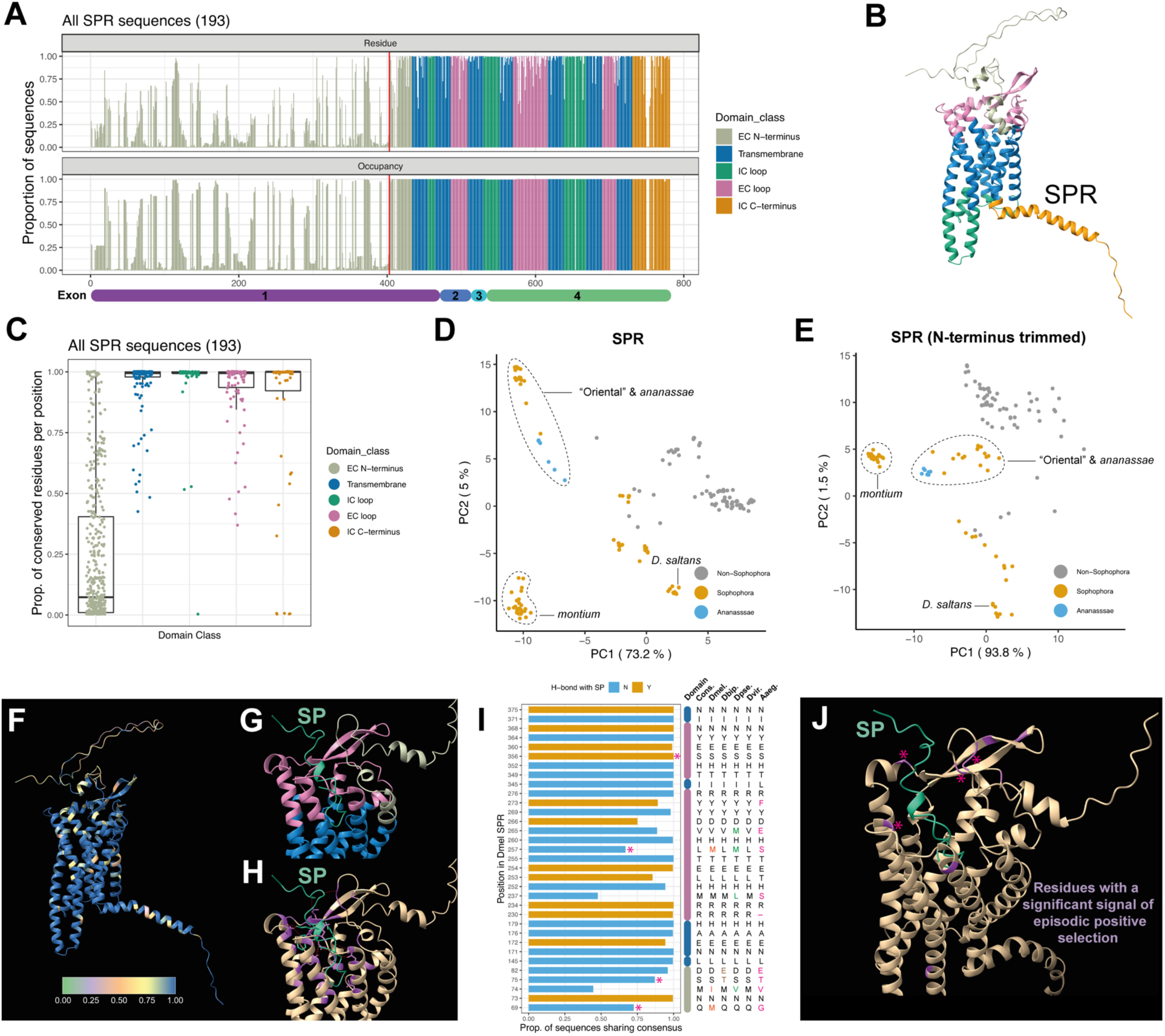
SPR residues showing evidence of episodic positive selection are enriched in the ligand-facing domains. (A) A consensus sequence based on MAFFT alignment of the resolvable amino acid sequences of *SPR* coding sequences. The top plot gives the proportion of sequences with the consensus amino acid in the same position, while the bottom plot gives the proportion of sequences in which each position is occupied in the alignment. Each residue is coloured based on the functional domain to which it belongs based on the UniProt annotations. The red line indicates the start of the conserved region we use in the molecular evolution analyses. Beneath the plot, we show the corresponding exon that encodes each consensus residue. EC = extracellular, IC = intracellular (B) The AlphaFold prediction of the structure of *D. melanogaster* SPR as downloaded from UniProt (AF-Q8SWR3-F1) and coloured by the domain each residue belongs to based on positions listed in the UniProt ‘Features’ table. (C) A boxplot showing the proportion of residues at each position that matched the consensus residue (*i.e.,* the degree of conservation at each position). Residues are plotted separately according to their domain class of origin. (D) PCA plot based on BLOSUM62 substitution scores from the MAFFT-aligned SPR protein sequences. The percentage values in the axis titles reflect the proportion of variance explained by a given PC. Points are coloured based on whether they correspond to *Sophophora-Lordiphosa,* non-*Sophophora-Lordiphosa,* or *ananassae* subgroup species. (E) As (D) but after removing the non-conserved region of the N-terminus (*i.e.,* the region preceding the red line in (A)). (F) The same prediction shown in (B) but with residues coloured by the proportion of conserved residues per position (see also Supplementary movie 1). High values indicate high conservation. (G) The ColabFold top-ranked prediction of the interactions between SP (shown in green) and SPR (residues coloured by domain). (H) As in (G) but with predicted contact residues coloured purple and predicted hydrogen bonds between SP and SPR residues shown with red-dotted lines (See also Supplementary movies 2 and 3). (I) A bar chart showing the proportion of sequences sharing the consensus residue for each predicted contact residue. Bars are coloured by whether the residue is also predicted to form a hydrogen bond with SP. Asterisks denote predicted contact residues for which we detected evidence of episodic positive selection using *MEME.* Alongside the plot, coloured bars, using the same colour scale as in (A-C), denote the functional domain the residue falls within. The two adjacent blue bars denote separate, consecutive transmembrane domains. Alongside are the corresponding amino acid residues in each position for each of the consensus (‘Cons.’), *D. melanogaster* (‘Dmel.’), *D. bipectinata* (‘Dbip.’), *D. pseudoobscura* (‘Dpse.’), *D. virilis* (‘Dvir.’), and *A. aegypti* (‘Aaeg.’) sequences. Residues that depart from the mode among these plotted sequences are coloured. (J) As (G) but colouring only the 10 residues in SPR for which we detected evidence of positive selection using *MEME*. Asterisks denote predicted contact residues that show evidence of positive selection.

### SPR sites with evidence of episodic positive selection are disproportionately located in predicted extracellular facing domains

Several residues in the extracellular loops and extracellular facing transmembrane domains – regions likely to be critical for ligand-binding – showed reduced conservation (Figure 6A, F; Supplementary movie 1). To assess whether these sites are under selection, we used *FUBAR* (60) to test for evidence of pervasive diversifying selection at individual sites in the N-terminus trimmed SPR sequence. We detected evidence of *pervasive* diversifying selection at 0/377 sites and purifying selection at 357/377. We followed this analysis with a test for *episodic* positive selection at individual sites, implemented through *MEME* (61). In this analysis, we detected evidence of episodic positive selection at 10/377 sites. Running the same analysis separately for the *Sophophora-Lordiphosa* (91 species) and non-*Sophophora-Lordiphosa* species (89 species), we identified 4/371 and 5/376 sites respectively as showing evidence of episodic positive selection (the identity of these positively selected sites did not overlap between the two analyses). Therefore, we found no evidence that the proportion of sites experiencing episodic positive selection was elevated in the *Sophophora*, which includes the lineages in which *SP* showed greatest evolutionary change.

Intriguingly, of the 10 positively selected sites identified in the phylogeny-wide analysis, 9 fell within extracellular domains: 3 in the N-terminus region close to the start of the first transmembrane domain, 6 across the three extracellular loops, and then one in the third intracellular loop. Using ColabFold (62–65) to generate a model of SP-SPR interactions (*SI Appendix, Fig. S14A-C*) and ChimeraX (66), we detected 33 residues in SPR that were predicted to interface with SP, of which 12 were additionally predicted to form hydrogen bonds with SP residues (Figure 6G-I; Supplementary movies 2, 3). Most of these residues were highly conserved across the 193 sequences: 12/33 were 100% conserved and 20/33 were >99% conserved (Figure 6I). But this level of conservation was not atypical among the extracellular facing residues: those that neighboured the predicted contact residues were similarly well conserved (*SI Appendix, Fig. S14E*). Of the 13 less well conserved residues, none showed clear evidence of concerted change among *ananassae* subgroup, Oriental lineage, or *Sophophora* species. However, there was a significant enrichment of sites showing significant evidence of episodic positive selection among the 33 predicted contact residues (4/33; *χ^2^*=12.56, *df* =1, *p*=0.0004; marked by asterisks in Figure 6I,J). Overall, therefore, while we do detect evidence that sites in the putative SP-binding pocket of SPR have undergone episodic positive selection, the number of changes does not appear to be elevated in the *Sophophora-Lordiphosa* radiation where the major genomic and functional changes (e.g., 25) in SP have occurred.

### A validated mutational route that SPR could take to decouple responses to SP and MIPs remains unexploited in drosophilids

If receipt of SP is associated with a net reduction in female fitness, the potentially deleterious effects of disrupting MIP-SPR interactions may constrain SPR’s ability to evolve to defend against SP binding. However, substitution of certain residues in SPR can have decoupled effects on the receptor’s sensitivity to its different ligands. Specifically, replacing the QRY motif at the boundary between the second intracellular loop and third transmembrane domain with the DRY motif more widely found in class A GPCRs is associated with a decrease in the responsiveness of SPR to SP, but not the ancestral MIP ligands, in *in vitro* assays (51). Yet in no drosophilid did we detect change at this position. Therefore, this potential avenue through which substitution of a single amino acid, albeit requiring change at two nucleotide positions, could reduce sensitivity to SP without affecting pre-existing ligand interactions remains unexploited.

## Discussion

Over the past few decades, we’ve built up a detailed understanding of the function of SP in *D. melanogaster*. We know that it is required for the normal assembly and disassembly of seminal storage and trafficking structures (“microcarriers”, 28); that it triggers an extensive range of physiological and behavioural changes in females, at least some of which are mediated by its interactions with SPR in female reproductive tract neurons (34–36); and that SP’s effects in females are extended via its binding to the surface of sperm, a process facilitated by a network of other male-derived proteins (29, 30, 32, 33). And yet, previous work has suggested that despite its integral roles in *D. melanogaster* reproduction, and despite the complex sperm-binding machinery with which it interacts, *SP* is restricted to drosophilids and perhaps, therefore, a drosophilid innovation (48). Consequently, *SP* represents a powerful system in which to chart the origin and diversification of function in a novel gene across different lineages. The cross-species analysis of *SP* and *SPR* genes that we report here makes several contributions to this.

The first relates to the phylogenetic distribution of *SP* genes. Previous work traced *SP* as far back as the split between *D. melanogaster* and *D. virilis* and identified three apparently independent loss events in the non-*Sophophora* species *D. grimshawi, D. mojavensis,* and *D. albomicans* (25, 48, 49). The data we present here pushes the origin of *SP* back to the base of the *Drosophilinae.* We also showed that *SP* is present in a genome of *D. albomicans* and that the losses in *D. grimshawi* and *D. mojavensis* are not species-specific, but instead cover much larger radiations, including all 55 members of the Hawaiian radiation that we sampled and, independently, the lineage leading to the *annulimana*, *bromeliae*, *nannoptera,* and *repleta* groups. Alongside, we detected evidence of additional losses in *H. duncani* and a lineage of *Scaptomyza*. Given the critical role of *SP* in many aspects of *D. melanogaster* reproduction these losses are intriguing and may reflect a lower functional importance of *SP* outside of the *Sophophora*. Consistent with this, SP injection experiments suggest that the ability of SP to reduce female sexual receptivity is restricted to the *melanogaster* group (25). Where *SP* has been lost it may be that its functions have been taken over by non-homologous proteins. Indeed, there is a clear precedent for non-homologous reproductive proteins being used to achieve similar phenotypic endpoints in different insect species (20, 21).

The second is our detection of substantial variation in copy number and sequence in the *Sophophora-Lordiphosa* radiation, a feature that was particularly pronounced in the “Oriental” lineage and *ananassae* subgroup. Outside of the *Sophophora-Lordiphosa, SP* is almost invariably a single (or 0) copy gene. But inside, where we detect repeated, independent duplication of *SP* across lineages, the story is quite different. Repeated duplication is at its most extreme in the *ananassae* subgroup, where we see as many as 7 copies present in *D. pseudoananassae nigrens.* Importantly, in this subgroup the intraspecific paralogs are not identical in sequence. Instead, they generally showed considerable variation in length and amino acid composition. What, then, are the functional consequences of this intraspecific diversity? If these paralogs are all interacting with SPR and varying to different degrees in their C-terminal sequences, do they vary in the efficiency with which they bind SPR? Or are they specialised for different receptors? We know, for example, that *D. melanogaster* SP can activate another GPCR, Methusaleh (Mth*), in vitro.* However, SP-Mth interactions don’t seem to be required for the post-mating increase in egg production or reduction in sexual receptivity – at least in *D. melanogaster* (67). There is also evidence that SP can induce some of its effects in *D. melanogaster* independently of *SPR,* pointing to the potential existence of additional, unidentified receptors (68). Thus, it’s possible that SP might be evolving in some lineages to make use of a wider set of receptors, a process that copy number amplification might facilitate.

But what about the other regions of SP proteins, the regions beyond the post-mating response stimulating C-terminus? After all, it’s the N-terminus and midsection regions that we show to be most variable between homologs. In *D. melanogaster*, we know that SP binds to sperm at its N-terminus; the N-terminal WEWPWNR motif then remains bound to SP after the rest of the peptide is cleaved (33). Our data suggest that clear variants of this motif are restricted to at least one SP copy in the Muller D2 neighbourhood of each member of the “Oriental” lineage (except for *D. oshimai*). This raises two possibilities: (1) the ability of SP to bind sperm is restricted to SP copies carrying variants of this motif in this lineage; (2) the protein network that underlies SP-sperm binding can facilitate attachment using a wide set of N-terminus sequences. Such flexibility might stem from rapid evolution of the sequence or identity of SP network proteins, or of any sperm surface proteins that SP might interact with. Indeed, there is evidence that several sex peptide network proteins have experienced recurrent positive selection in the *melanogaster* group (49). Flexible use of N-terminus sequences may also be due to some inherent, accommodating property of the sperm binding apparatus. Consistent with this, no such WEWPWNR motif is present in *D. melanogaster* Dup99b, but it nevertheless binds to sperm, albeit only to the sperm head and only during the first few hours after mating, unlike SP (69). But whether Dup99b is relying on the same network of proteins as SP to bind to sperm, or perhaps interacting with distinct proteins on the sperm surface, remains untested. Where multiple, divergent SP copies are present, as in the *ananassae* subgroup for example, we may be seeing specialisation in the N-terminus region that relates to a given peptide’s mechanism of interacting with the female: while some may be adapted to bind sperm, others might be adapted to enter into the hemolymph, as *D. melanogaster* SP has been shown to do (70).

The third relates to the genomic distribution of *SP* genes. We observed that *SP* genes have frequently translocated to new genomic locations, a feature that’s particularly pervasive in *Sophophora.* A key question here is to what extent these translocations have shaped the evolution of SP function. For example, did the ‘Muller D1’ to ‘Muller D2’ translocation at the base of the *Sophophora-Lordiphosa* lineage open the door to the novel evolutionary trajectories taken within this lineage? We could imagine that this translocation placed *SP* within a new *cis*-regulatory environment that changed either the strength, timing, or tissue-specificity of its expression, thereby exposing it to new selective forces. Our data suggest against this at a global level, at least in relation to tissue-specificity, as we observed accessory gland biased expression of *SP* in *D. virilis*, a species that pre-dates the translocation. However, it’s possible that subsequent translocations within some branches of the *Sophophora-Lordiphosa* radiation are associated with shifts in expression pattern to new subregions of the male reproductive tract. After all, the translocated Muller element E copy of *SP* in *D. melanogaster* (*Dup99b*) is expressed not in the accessory glands, like *SP,* but in the ejaculatory duct (44).

The fourth is our failure to detect a clear association between evolutionary change in *SP* and microcarriers. The presence of LipidTox^+^ microcarriers in *S. flava,* which lacks a copy of *SP*, suggests that *SP* isn’t necessary for microcarriers. Moreover, the absence of canonical LipidTox^+^ microcarriers in *SP*-encoding species, such as *D. immigrans*, suggests that a copy of *SP* isn’t sufficient. Thus, SP playing a role in structuring microcarriers might itself be a derived trait, as might its association with microcarriers more broadly (*e.g.,* as a microcarrier cargo), thereby casting doubt on the idea that this could be SP’s ancestral function (15). This association might be relatively recent, as even among many *SP*-bearing *Sophophora* species variation in *SP* copy number and sequence doesn’t seem to be obviously connected to variation in microcarrier morphology. Our stainings also raise the issue of what exactly defines a ‘microcarrier’. The detection of LipidTox^-^, ellipsoid, microcarrier-like structures and LipidTox^-^flocculence raises the question of whether these represent fundamentally different structures to microcarriers or whether taking up LipidTox (an indicator that they contain large quantities of triglycerides and other nonpolar lipids (28)) is a feature of some, but not all, microcarriers – a feature that our data suggest appears to have been gained and lost repeatedly, perhaps in line with higher-level dietary or metabolic changes.

Our data point to lineage-accelerated evolution of *SP* within the *Sophophora-Lordiphosa* radiation, marked by repeated, independent rounds of gene family expansion. What forces are driving this trend? And why does it appear to be so much more pervasive in these lineages? A tempting response to the latter question is that it coincides with some evolutionary shift in the activity of SP. A previous study of 11 drosophilid species found that the ability of conspecific SP to reduce female sexual receptivity is confined to the *melanogaster* group (25). Intriguingly, this gain in responsiveness to SP also appears to coincide with both the gain of robust expression of *SPR* within the female reproductive tract and the ability of SP (derived from *D. melanogaster*) to bind to female reproductive tract tissue (25). Viewed from a sexual conflict perspective, therefore, we might be detecting the effects of a sexually antagonistic coevolutionary arms race that was initiated after the acquisition of new functions by SP. If responding to SP is deleterious to female fitness (*e.g.,* 43), then females might evolve resistance, in turn selecting for structurally divergent copies of *SP* through which males can overcome that resistance. Indeed, the numerous, structurally divergent copies of *SP* expressed by members of the *ananassae* subgroup are consistent with theory suggesting that males might gain from transferring diverse sexually antagonistic seminal fluid products simultaneously as a ‘combination’ strategy to overcome the evolution of resistance (71).

But if females are evolving resistance, then we fail to find strong evidence that it is occurring through *SPR*. The only region of SPR where we see extensive evolutionary change is in the N-terminus. But there are reasons to believe that any role this region plays in ligand binding is relatively minor (see *SI Appendix, Supp. Text*). And while we detect evidence of episodic, but not pervasive, positive selection at 10 sites in SPR beyond the N-terminus, these changes were distributed throughout the phylogeny and without enrichment in the lineages experiencing accelerated evolution of *SP.* Why, then, are we not seeing SPR rapidly evolving in concert with SP? One explanation is that the evolution of SPR is constrained by the need to maintain interactions with additional ligands, such as MIPs. But we know from *in vitro* assays that there is at least one mutational route for SPR that has decoupled effects on MIP and SP binding, negatively affecting the latter more strongly than the former (51). And yet across 193 drosophilid SPR sequences we failed to find any instance where this route has been exploited (though it would require a double mutation). Another possibility is that any resistance to SP-mediated sexual antagonism might be mediated up- or down-stream of SP-SPR binding, such as via mechanisms that degrade SP, block it from binding, or in how the neural circuitry that SPR feeds into responds to SP. One such potential action point is sperm cleavage: if females could block the cleavage of SP from the surface of the sperm – the mechanism for which remains uncharacterised – then they could markedly reduce the timeframe over which SP’s effects are active (∼10-14 days in D. melanogaster; 30).

But limited evolutionary change in *SPR* is also consistent with another possibility: the role of sexual conflict in driving the evolution of SP might be relatively weak. While the functions of SP are often framed in terms of male manipulation of female reproductive decision-making or collateral damage in the pursuit of improved performance in sperm competition (e.g., 72–75), the evidence base for this is not strong (reviewed in 15). There is theoretical support for antagonistic effects of seminal proteins in general (*e.g.,* 83) and empirical support in *D. melanogaster* for the antagonistic effects of seminal proteins (77) and SP specifically (45, 78), but there is also empirical support for positive, neutral, and context-dependent (*i.e.,* in relation to female nutritional state) effects of SP on female fitness (46, 47, 79), as well as considerable uncertainty over the extent to which fitness measurements made in these laboratory studies reflect those experienced by wild-living populations (15, 80). Indeed, the recently demonstrated gain of both robust *SPR* expression in a subset of female reproductive tract neurons and the ability of SP to bind to reproductive tract tissue in the *melanogaster* group (25) suggests that females did, at least at some point, benefit from responding to SP, perhaps as part of a mechanism through which the receipt of sperm could be aligned with the induction of reproductive processes. Whether that may have subsequently initiated conflict – and what the associated genomic consequences might have been – remains to be resolved.

The high diversity we detect in SP coupled with the low diversity documented in the receptor mirrors patterns that have been recorded at the morphological level. While there are instances of correlated diversification between male-specific traits and female defensive traits that are suggestive of coevolutionary arms races over mating rate, such as in the clasping and anti-clasping structure of *Gerris* water striders (81, 82), there are also a great many male-specific traits across arthropods that show rapid interspecific diversification without apparent change in the female structures they contact (83). In such cases, the diversification of male traits has traditionally been thought to be driven by sexual selection under female choice (17). Others, notably Eberhard and Cordero, have argued that seminal proteins may similarly be evolving under female choice (16, 18). In terms of SP, this may mean that a female gains from discriminating between different males on the basis of the dose or sequence of SP she receives, tailoring her use of sperm or overall reproductive investment in response (15, 84). This, in turn, may select for diversification in SP sequence or increases in expression (perhaps via copy number amplification) to meet female preferences (85).

Ultimately, sexual conflict and sexual selection by female choice needn’t act entirely independently (15, 86, 87). But disentangling the historical importance of these forces in driving the evolution of SP-female interactions continues to present a major challenge, just as it has for many other traits since work in this area began (83, 84). Our belief is that by charting the origin and diversification of SP-female interactions at a mechanistic level, we may be able to draw more robust inferences regarding the evolutionary forces that are shaping this recently evolved system in *Drosophilinae*.

## Methods

Detailed sample information and accession numbers for the 271 genomes used in this study are provided at https://osf.io/24unk. SP sequences were identified by combining reciprocal blast of the *D. melanogaster* SP C-terminus sequence with protein sequence, gene structure, and synteny analysis. Signal peptides were removed from the translated protein sequences using SignalP-6.0 run on ‘fast’ mode and specifying ‘Eukarya’ as the organism (88). For the microcarrier stainings, accessory glands were stained in 100μl of 1:50 LipidTox Deep Red Neutral Lipid Stain (Invitrogen, H34477) in PBS for 1 hour with 1μl 1:100 DAPI added for the last 15 mins. Tests for selection in SPR coding sequence were conducted using Datamonkey (89, 90). All code, extracted coding and protein sequences, expression data, and protein models are available at https://osf.io/tzu6v/. A detailed methods section is available in the *SI Appendix*.

## Acknowledgements

We thank the reviewers and editor for thoughtful and valuable feedback on earlier drafts of this manuscript; Clive Wilson and Mark Wainwright for helpful feedback on the microcarrier part of this work; Julianne Peláez and Noah Whiteman for providing us with *Scaptomyza* stocks; and Barbara Parker, Kathy Darragh, Pavitra Muralidhar, and Carl Veller for valuable discussion. This work was supported by a long-term fellowship from the Human Frontier Science Program Organization (LT000123/2020-L to B.R.H.), a UC Davis College of Biological Sciences Dean’s Circle Summer Undergraduate Research Program Fellowship to A. A-H., and National Institutes of Health F32 GM135998 to B.Y.K., R37-HD038921 to Mariana Wolfner, and R35 GM122592 to A.K.

