## Supplementary material for "Decoupled evolution of the *Sex Peptide* gene family and *Sex Peptide Receptor* in *Drosophilidae*": SI Appendix

**This PDF file includes:**

Supporting text
Figures S1 to S15
Legends for Movies S1 to S4
SI References

**Other supporting materials for this manuscript include the following:**

Movies S1 to S4

### Supporting Text

The only region of SPR where we see extensive evolutionary change is in the N-terminus. But there are reasons to believe that any role this region plays in ligand binding is relatively minor. For one, the only predicted contact residues within the N-terminus that we identified were all directly adjacent to the transmembrane domain and fell within the region that showed highest conservation. A counter to our failure to detect contact residues in the faster evolving regions of the N-terminus would be to point out that the 3D structure of the N-terminus was predicted with the lowest confidence of any part of the protein (*SI Appendix, Fig. S14A*), and we may therefore be missing potential interaction sites with SP. That being said, the low confidence of this region's structure is also consistent with it being intrinsically disordered, a feature known to be present in the N-termini of other GPCRs such as human beta-2 adrenergic receptor (1). This disorder may reflect a lack of functional importance for ligand binding and evolution under relaxed selection. Indeed, the N-termini of other GPCRs appear to be considerably more mutationally tolerant than other parts of the protein (2). However, there are examples, such as rhodopsin, where the N-terminus plays a key role in structuring the region surrounding the ligand-binding site (1). These broader features of GPCRs aside, the best evidence we have against the N-terminus of SPR being a strong driver of responsiveness to SP comes from previous domain swapping work, where different domains of *D. melanogaster* SPR were swapped with the homologous regions from *A. aegypti* SPR and the effects on SP binding tested using *in vitro* assays. The N-terminus swap, which extended deep into the first transmembrane domain, conducted as part of this previous work had a much more modest effect on SP binding relative to swapping regions containing transmembrane and extracellular loop domains (3).

### Materials and methods:

***Genomes.*** 271 genome sequences for 264 species (with 2-3 genomes from different strains for 6 species) were obtained in the same manner as described in previous work (4, 5). We sequenced and *de novo* assembled genomes for 108 species (Kim et al., *in* prep), downloaded 155 publicly available assemblies from NCBI databases (4, 6–30), and downloaded unassembled reads from the NCBI Sequence Read Archive for 8 species (5, 14). Detailed sample information and accession numbers are provided at https://osf.io/24unk. We sequenced on Oxford Nanopore MinION and Illumina NovaSeq instruments and assembled *de novo* genomes with the hybrid approach described in Kim *et al.* (4) and Kim *et al.,* (in prep). Genomes of species for which only short read data were available were assembled with SPAdes 3.13.1 (31).

***Phylogeny.*** A species tree was generated from 250 orthologs, again following the workflows established in previous work (4, 5). Briefly, we identified single-copy dipteran orthologs using BUSCO v5 (32) and randomly selected 250 genes identified as single-copy in the assemblies and missing for no more than 10 species. Each ortholog was individually aligned with MAFFT (33) and gene trees were inferred with IQTREE 2.2.0.7 (34). A species tree was inferred from the gene trees with ASTRAL-III (35).

***Identifying SP genes***. To identify SP orthologs, we first searched each genome by tBLASTn using a liberal E-value cut-off of 100. As the query, we used the *D. melanogaster* C-terminus sequence (KWCRLNLGPAWGGRC), which is entirely encoded by the second exon. This is the best conserved domain between SP and Dup99b in *D. melanogaster* (36) and which previous work on SP sequences from 10 species had shown to be the only strongly conserved region (37). We extracted 350nts either side of the start and end points of each returned hit and reciprocally queried the extracted sequence by BLASTx against the *D. melanogaster* reference protein list, retaining only those sequences that gave an SP paralog (SP or Dup99B) as the top hit. The extracted sequence length is more than sufficient to capture the full *D. melanogaster SP* (288nts) and *Dup99b* (310nts) genes. To be included as an *SP* gene in the total species counts (*i.e.,* those given in Figures 1-4), a sequence had to meet these BLAST criteria and give a recognisable SP-like protein sequence when aligned to the *D. melanogaster SP* gene structure (*e.g.,* no premature stop codons). For species that returned no hits through this approach, we used two follow-up approaches (within each of the Hawaiian and *annulimana*, *bromeliae*, *nannoptera* and *repleta* ('ANBR') radiations, we chose 5 representative species). First, we used tBLASTn to query the full-length, including N-terminus, resolved SP protein sequences from a selection of 11-14 species from across our phylogeny against each apparently *SP*-less genome. In only one of 13 species tested using this approach (*D. neohypocausta*) did we detect a *SP* homolog. In the remaining 12 species we detected nothing that resembled a *SP* homolog, nor any individual hit that fell within the Muller D1 neighbourhood despite using a liberal E-value cut-off. Second, we repeated this approach using tBLASTn to query a selection of 6-9 full-length SP sequences from a phylogenetically varied set of species against only the Muller D2 neighbourhood of each of the remaining 12 species. In no case did we detect evidence of any remaining fragment of *SP.*

***Mapping the syntenic neighbourhoods of SP genes.*** To position *SP* hits within their chromosomal context, we sought to identify and reconstruct syntenic gene neighbourhoods surrounding each *SP* hit. To do this, we first used the NCBI genome viewer to identify and extract the protein sequences of genes immediately up and downstream of *SP* in the *D. melanogaster* reference genome (DmelRS6). In chromosomal order, these genes were *tartan, CG33262, snky, capricious, Sfp70A4, CG42481, CG43147, (SP), CG14113, CG17687, Nplp2,* and *CG14111.* We then searched each of these sequences against each species' genome using tBLASTn, extracting the coordinates for the top returned hit in each case. Of these genes, we focused on *capricious, CG17687,* and *CG14111* as these were particularly well-conserved. In a minority of cases, contig breakpoints fell within a neighbourhood preventing us from fully reconstructing the region. In one case (*D. ananassae*), we were able to stitch two contigs together that spanned an *SP*-containing neighbourhood by cross referencing to the *D. ananassae* reference genome (DanaRS2.1). In another (*L. mommai*), we used the other *Lordiphosa* species to stitch two Muller D2 contigs together that separated *capricious* from *SP, CG14111,* and *CG17687,* allowing us to plot the position of *SP* in Figure 3. In a small number of cases, this search revealed the presence of *SP* genes within the syntenic gene neighbourhood that hadn't passed our reciprocal BLASTx test. These were *D. nasuta, D. immigrans, D. immigrans (kari17)*, *D. guttifera, D. sulfurigaster bilimbata, D. seguyi,* and *D. oshimai*. We included these for further analysis based on their shared chromosomal location with *SP* genes that did pass the reciprocal blast test in other species and the SP-like protein sequences that we were able to resolve for them.

For SP hits left unlocated by this approach, we first repeated this same process for the gene neighbourhood surrounding *Dup99b* (*CG34296, dmrt99b, gycalpha99b*). If that failed to identify their position, we extracted variable, larger regions of sequence surrounding the hit on each *SP*-containing contig and used a combination of BLASTn and tBLASTn searches against the *D. melanogaster* reference genome. From this, we identified syntenic gene neighbourhoods, extracted the protein sequences for these *D. melanogaster* genes from Flybase (38), and repeated the approach used for *SP* and *Dup99b* with these new neighbourhood genes. This analysis showed that in almost all species SP genes were located within a small number of highly conserved syntenic neighbourhoods. Neighbourhood plots were constructed in R (V4.1.1) using bespoke scripts that used ape (39), ggtree(40), and ggplot2 (41).

***Reconstructing SP gene structure and protein sequence.*** To annotate *SP* gene structure, we generated a MAFFT alignment (42) of all our identified *SP* sequences and identified landmarks (start and stop codons, exon/intron boundaries) that were conserved with the *D. melanogaster SP* and *Dup99b* sequences. This approach was previously used by Tsuda *et al.* (37) on a smaller bank of 10 *SP* sequences. As in Tsuda *et al.* (37)*,* our alignment revealed broad conservation of both the start and stop codons and the intron donor (consensus: GTAAGT) and acceptor (AG) sequences across species. In the small number of sequences where these landmarks were not present at the conserved position in our alignment, we selected the closest plausible option to the consensus sequence position that was conserved among similar species. We further required that the selected option had to maintain the KWCRLNLGPAWGGRC-like reading frame in exon 2. Of 269 identified SP-like nucleotide sequences, we could resolve SP-like protein sequences for all but 34, most of which fell outside syntenic gene neighbourhoods that housed *SP* in other species, suggesting that they are unlikely to be real *SP* genes. Signal peptides were removed from the translated protein sequences using SignalP-6.0 run on 'fast' mode and specifying 'Eukarya' as the organism (43). A signal peptide sequence was detected with a probability of >0.9992 for all but four sequences, two of which came from the *montium* subgroup species *D. punjabiensis* (0.0584 and 0.0055) and the other two from its sister species in our dataset *D. watanabei* (0.1284 and 0). In these cases, it might be that these represent non-homologous sequences that bear a resemblance to SP in the C-terminus or that these SP proteins are no longer secreted into the extracellular environment. At 34aa, these were the shortest SP orthologs we detected. The longest we detected were a pair of 76aa paralogs (pre-cleavage) in the non-*Sophophora* species *S. latifasciaeformis.*

Our results showed strong conservation of the C-terminus motif between sequences. While this may suggest that this C-terminus sequence is the defining characteristic of SP family proteins, the high degree of conservation we detect might be an artefact of us ourselves having used this sequence as the defining characteristic of an *SP* gene. Because of this search design, while we can readily pick up SP sequences with highly diverged N-termini, we would not be able to detect an SP sequence with a conserved N-terminus and highly diverged C-terminus. What we can conclude, however, is that the C-terminus appears highly conserved, that certain residues within the C-terminus are almost invariable, and that where *SP* genes are detected based on the C-terminus sequence the N-terminus and midsection regions appear to be considerably more variable in sequence.

***Expression analysis.*** For 19 species, we identified RNA-seq datasets on NCBI that we could use to test for expression of our predicted SP orthologs. For each of these species, we used BLASTn to search each ortholog against a given species' genome. We then used the graphics option in the NCBI blast suite to view the hit mapped to the genome, extracted the associated GeneID and gene class identity, and loaded available RNA-seq exon coverage tracks. For each loaded track, we recorded the associated metadata (tissue, sex, dataset accession, *etc.)* and the peak exon coverage within the SP ortholog coding sequence. For each RNA-seq sample, we extracted the equivalent value at the housekeeping gene *RpL32* locus. We calculated relative expression values by adding 0.1 to the peak *SP* exon coverage value and dividing it by the equivalent *RpL32* value. We further used each *SP* hit in each species for which we had RNA-seq data to confirm our correct annotation of exon/intron boundaries.

During this analysis, we observed that many of the *SP* genes that we detect are incorrectly annotated in the species' reference genomes. In the case of *D. bipectinata,* 4 of its *SP* genes are designated as 'uncharacterised long non-coding RNAs' (LOC108127578, LOC108127576, LOC108127577, and LOC108127621). This was a common designation for *SP* genes, one that we also saw for *D. kikkawai* (LOC108075137), *D. takahashii* (LOC108063118 and LOC108063123, but LOC108062332 was listed as a *SP* homolog), *D. ananassae* (LOC116655869 and LOC26513946), *D. willistoni* (LOC124460714), *D. virilis* (LOC26531923), and *D. ficusphila* (LOC108095416 and LOC108100235, but LOC108095478 was listed as a *SP* homolog). Some *SP* homologs that we detected in each of *D. biarmipes and D. ficusphila* showed clear male-biased expression but were unaccompanied by any gene annotation, not even a long non-coding RNA (*SI Appendix, Fig. S15*). Thus, automated genome annotation pipelines appear to often fail to recognise the homology of *SP* genes and, in some cases, fail to recognise them as genes entirely. These issues likely stem from a combination of the small size of *SP* family genes – a feature known to cause problems for gene finding tools (44, 45) – and it's low conservation beyond the C-terminus.

**SP *protein sequence analysis using PCA.*** PCA was performed using Jalview (v2.11.2.5) on a BLOSUM62 substitution matrix of all 229 resolved, signal peptide cleaved SP protein sequences from species shown in Figures 2 and 3 (46). Thus, the short paralogs in *D. watanabei* and *D. punjabiensis* for which we were unable to detect a signal peptide were not included. The transformed values were extracted from JalView and plotted using the gpplot2 package in R (v4.1.1).

***Constructing a* SP *protein tree.*** We inferred the evolutionary history of Sex Peptide proteins applying the Maximum Likelihood method with a Jones-Taylor-Thornton (JTT) matrix-based model (47) to a MUSCLE alignment of 233 extracted SP amino acid sequences from species shown in Figures 2 and 3. The tree with the highest log likelihood (-10372.14) is presented in *SI Appendix, Fig. S4*. Initial tree(s) for the heuristic search were obtained automatically by applying Neighbour-Join and BioNJ algorithms to a matrix of pairwise distances estimated using the JTT model, and then selecting the topology with superior log likelihood value. This analysis was conducted in MEGA X (48, 49).

***Fly rearing.*** The following strains were used for microcarrier stainings: *D. melanogaster* (RAL-517; 50), *D. malerkotliana malerkotliana* (mal0-isoC), *D. malerkotliana pallens* (Q120-isoG), *D. pseudoananassae pseudoananassae* (wau125), *D. pseudoananassae nigrens* (VT04-31), *D. varians* (CKM15-L1), *D. ercepea* (aag001 copy2), *D. bipectinata* (14024-038.07 nanopore), *D. ananassae* (OGS-98K1), *D. biarmipes* (361.3-iso11 1-11) *D. ficusphila* (iso1 L.10), *D. kikkawai* (4_2_3_2_1_2_1_1), *D. immigrans* (15111.1731.12), *D. rufa* (EH091 isoC L-3), *D. carrolli* (KB866), and *D. takahashii* (14022-0311.14). For most species, virgin males were collected within 8 hours of eclosion, held in groups of 7-10 individuals, and aged 8-10 days before their dissection and staining. The exceptions were *D. melanogaster*, which were used at 3-5 days (as in 51); *D. carrolli*, whose virgin status was unknown but which had been isolated from females 48h prior to dissection to allow accessory gland replenishment; and the *Scaptomyza* species, adults of which were generously provided by Julianne Peláez and Noah Whiteman. *S. pallida* and *S. hsui* were maintained on standard Bloomington media with thawed frozen chopped spinach. Non-virgin males from these species aged 4-6 days in *S. pallida*, 4-8 days in *S. hsui*, and 1-4 days for *S. flava* were used.

***Microcarrier stainings.*** Male reproductive tracts were dissected in 1x PBS in glass wells. Prior to staining, the PBS was removed from the wells and the tracts fixed for 30 mins in 200µl of 4% PFA (in PBS) solution on a rocker. The PFA solution was then removed, and the tracts subjected to three, 2-minute washes in 1x PBS on a rocker. After washing, we removed all excess tissue (including testes and seminal vesicles) except the accessory glands and pierced the glands with forceps to facilitate dye penetration (as in 51). Accessory glands were then stained in 100µl of 1:50 LipidTox Deep Red Neutral Lipid Stain (Invitrogen, H34477) in PBS for 1 hour with 1µl 1:100 DAPI added for the last 15 mins. During staining, the accessory glands were placed onto a rotator and covered to prevent exposure to light. After staining, the glands were washed 3 times in 1x PBS for 2 minutes each on the rotator. Finally, glands were mounted in 40µl Fluoromount 50 (Southernbiotech) on polylysine-coated slides and refrigerated until imaging. For each species, accessory glands from *D. melanogaster* were co-stained in the same well as a control to ensure successful staining. The species identity of mounted accessory glands was readily identifiable by morphological features of the accessory glands and microcarriers. At least 5 glands were mounted per species. Confocal images were taken using an Olympus FV1000 laser scanning confocal microscope.

***Identifying SPR orthologs***. To identify *SPR* orthologs, we used tBLASTn to separately query the coding sequence of each of the four *D. melanogaster SPR* exons against each species' genome. Each returned hit was then reciprocally blasted against the *D. melanogaster* reference protein list and all sequences that gave SPR as their top hit were extracted along with 1500 nucleotides up- and down- stream. The coordinates of each retained hit for each exon in each species were then extracted. We removed any sequences from one exon that were not contiguous with sequences from another and where all four contiguous exons were present (*e.g.,* an exon 1 hit on contig_A would be dropped if exons 1 through 4 were tandemly arranged on contig_B). All sequences from a given exon were then aligned using MAFFT and manually annotated based on features conserved with the *D. melanogaster SPR* sequence. The start codon of exon 1 aligned poorly at a phylogeny-wide level. In these cases, we selected the start codon closest to one that was conserved among closely related species that gave an SPR-like coding sequence. For 5 species, we were unable to resolve SPR sequences. In *Z. taronus*, a premature stop codon was present within the coding sequence of exon 4. For *S. montana* and *S. flava,* we failed to detect SPR-like sequences across either of its 4 exons when each *D. melanogaster SPR* exon was queried against the two species' genomes. In these species, the top hit for each exon gave a markedly lower score than the median across all species studied, suggesting loss of the whole protein-coding sequence (*SI Appendix, Fig. S13*). In the cases of *S. hsui* and *L. magnipectinata*, we found only 2 of the 4 exons gave a clear SPR match (*L. magnipectinata*: exons 1 and 4; *S. hsui:* exons 3 and 4), suggestive of a partial loss of coding sequence. Specific domains and their corresponding positions within the SPR protein sequence were extracted from the 'Features' section of the UniProt entry for SPR (Q8SWR3).

***Molecular evolution analysis.*** To test for selection in SPR coding sequence, we used a set of methods available through the 'Datamonkey' adaptive evolution server (52, 53). We used aBSREL (adaptive Branch-Site Random Effects Likelihood) to test if positive selection has occurred on a proportion of branches (54, 55), BUSTED (Branch-Site Unrestricted Statistical Test for Episodic Diversification) to test whether *SPR* has experienced positive selection at one or more sites on one more branches (56), FUBAR (Fast, Unconstrained Bayesian AppRoximation) to detect sites evolving under pervasive positive selection in a subset of branches (57), and MEME (Mixed Effects Model of Evolution) to detect sites evolving under episodic positive selection in a subset of branches (58). Each analysis was performed using default parameters on the conserved region (*i.e.,* N-terminus removed) of SPR sequences, aligned using a codon alignment generated using PAL2NAL (59) on the MAFFT-aligned protein sequences. For these analyses, the 193 sequences used to generate the consensus sequence plot was reduced to 182 to drop 3 sequences that were not in our phylogeny (*D. chauvacae*-like*, Leucophenga varia,* and *D. sp st01m*) and 8 where we had multiple lines per strain. A single strain was used for each of these species: *Z. indianus* (BS02)*, D. paulistorum* (14030-0771.06)*, D.immigrans* (15111.1731-12)*, D. robusta* (Kim *et al., in prep*)*, D. teissieri* (CT02)*,* and *D. willistoni* (14030-0811.00). Alongside the full species set, we repeated these analyses separately on alignments of the *Sophophora-Lordiphosa* and non-*Sophophora-Lordiphosa* species. A .json file for the results of each analysis is available at https://osf.io/tzu6v along with instructions of how to view the full output using https://vision.hyphy.org/.

***Binding site prediction.*** To predict the SP binding site, we ran the *D. melanogaster* SPR and cleaved SP sequences through ColabFold (https://colab.research.google.com/github/sokrypton/ColabFold/blob/main/AlphaFold2.ipynb). The cleaved form of SP was missing both the signal peptide and the first 7 residues of the mature peptide. Previous work has shown that SP is cleaved from the surface of sperm at a putative $R_7K_8$ trypsin cleavage site, with the remainder of the peptide left bound to the surface of the sperm (60). Additional work has shown that both the full length ($SP_{1\text{-}36}$) and N-terminally truncated form of SP ($SP_{8\text{-}36}$) are equally efficient in eliciting a receptor-mediated response in *in vitro* assays (61). However, we opted to use the cleaved form to match *in vivo* conditions. The top ranked model of SP-SPR interactions had the following metrics: local structural accuracy, pLDDT = 69.5; overall topological accuracy, pTM = 0.711; interface pTM = 0.812). Recent work has used a stringent cutoff of ipTM = 0.85 for calling high-confidence protein-protein interactions, observing that values between 0.55-0.85 perform better than random, with increasing accuracy at higher values (62). Thus, the prediction falls a little short of the high-confidence cutoff, but still suggests a reasonable level of confidence. As a further test, we compared the ipTM scores between *D. melanogaster* SP and each of the 44 members listed on Flybase of the Class A/rhodopsin-like G-Protein Coupled Receptors family to which SPR belongs. SPR gave the highest score of all (*SI Appendix, Fig. S14D*). The .pdb file of the top ranked model was then loaded in ChimeraX (v1.5)(63) and the 'interface residues' and 'hbond' commands used with default settings to identify putative contact and hydrogen bond forming residues in SPR, respectively. To compare ipTM scores between cleaved SP and other *D. melanogaster* Class A GPCRs, we extracted protein sequences from Flybase for each gene listed in the 'Class A GPCR neuropeptide and protein hormone receptor' gene group (FBgg0000195) and ran them through ColabFold (64–67) using default settings, as for SPR.

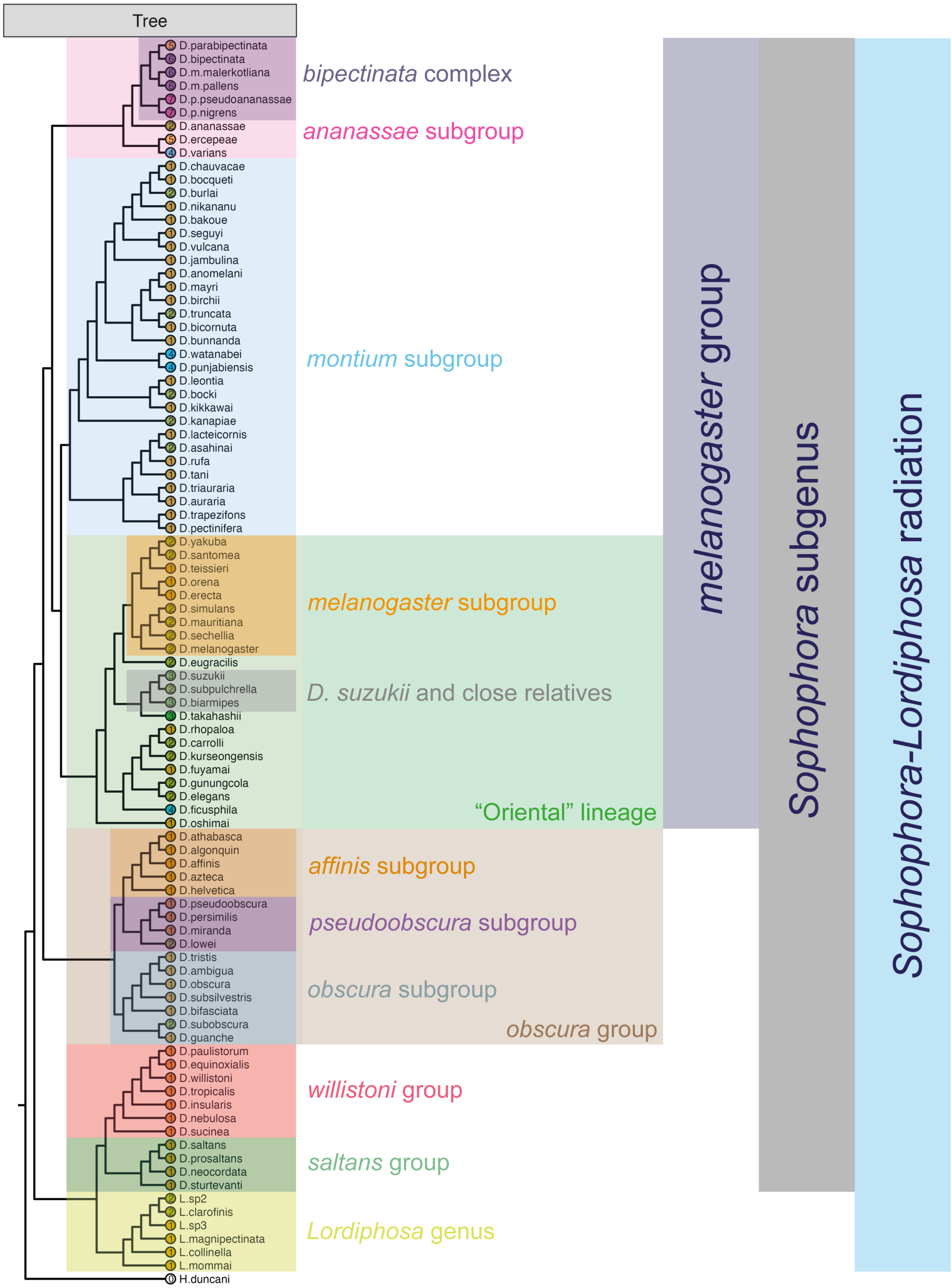


**Fig. S1. Taxonomic nomenclature used in this study.** This figure shows the phylogeny and *SP* copy numbers (given in coloured circles at each tip) from Figure 3 with branches coloured according to higher-order groupings we use in this study. The classification of the "Oriental" lineage follows the convention of Kopp and True (68). The positioning of *D. helvetica* within the *affinis* subgroup follows Gao *et al.* (69). A higher resolution figure can be downloaded at: https://osf.io/vcgbm.

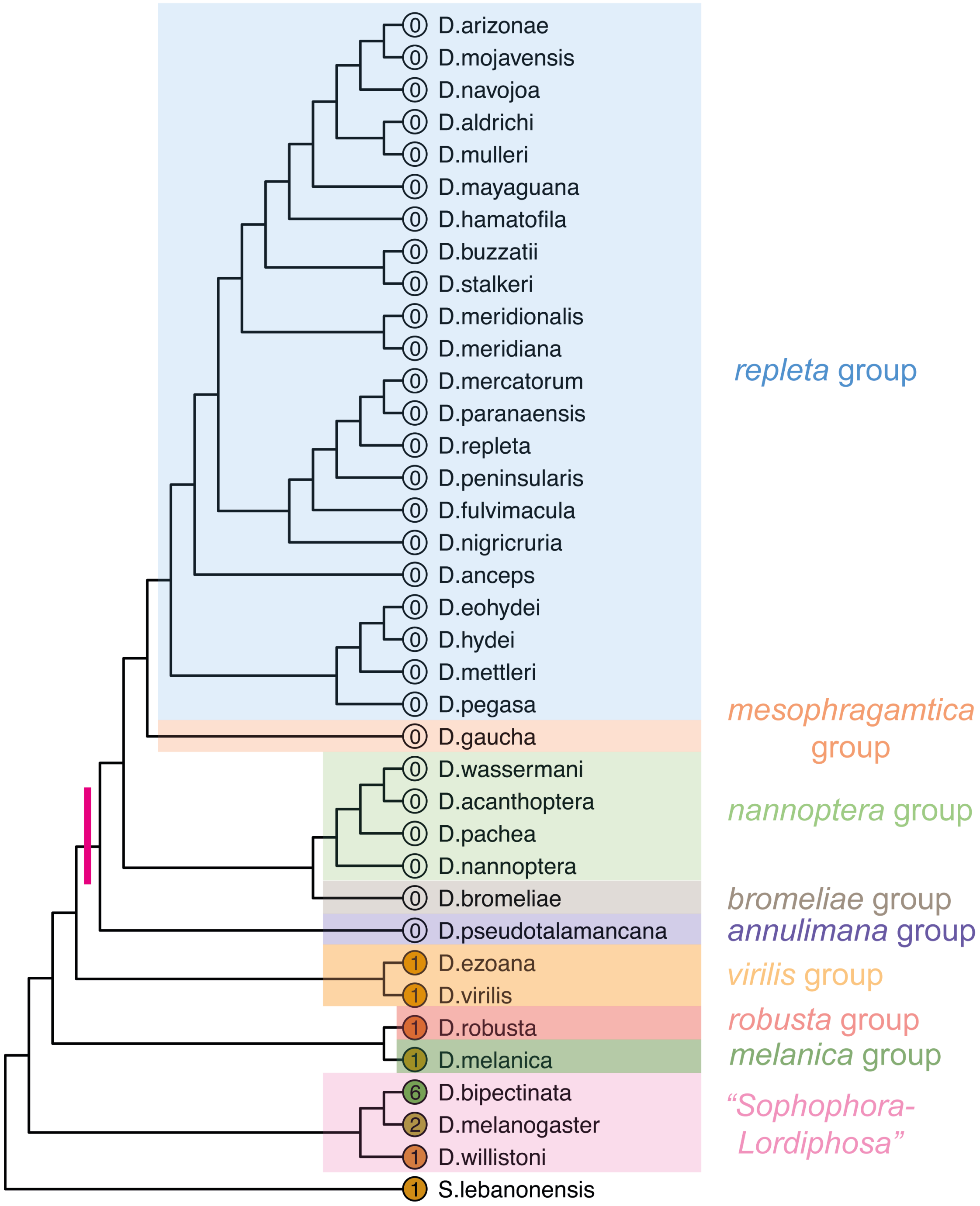


**Fig. S2. *SP* was lost at the base of the lineage containing the *annulimana, bromeliae, nannoptera, mesophragmatica*, and *repleta* groups.** This figure provides an expanded set of *repleta* group species relative to Figure 2. The number of SP genes detected is given in coloured circles at the tip of each branch. A pink bar denotes the branch on which we infer SP to have been lost. A higher resolution figure can be downloaded at: https://osf.io/qpem3.

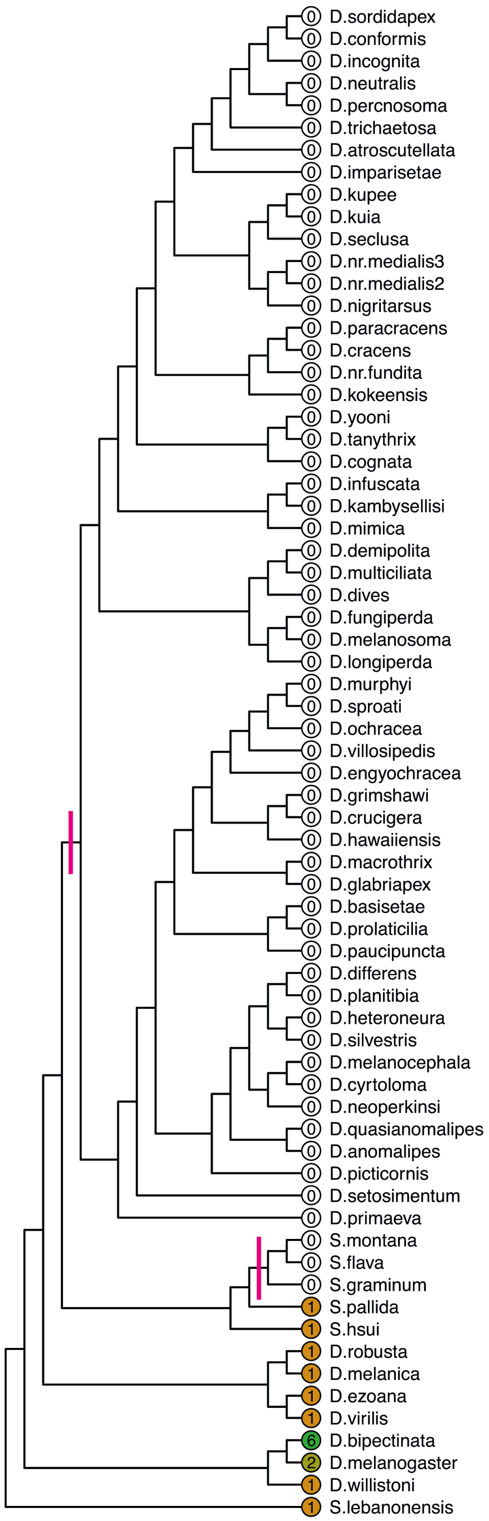


**Fig. S3. *SP* was lost in the Hawaiian group and a sublineage of *Scaptomyza*.** The number of SP genes detected is given in coloured circles at the tip of each branch. This figure provides an expanded set of Hawaiian group species relative to Figure 2**.** Pink bars denote the branches on which we infer SP to have been lost. A higher resolution figure can be downloaded at: https://osf.io/xbg57.

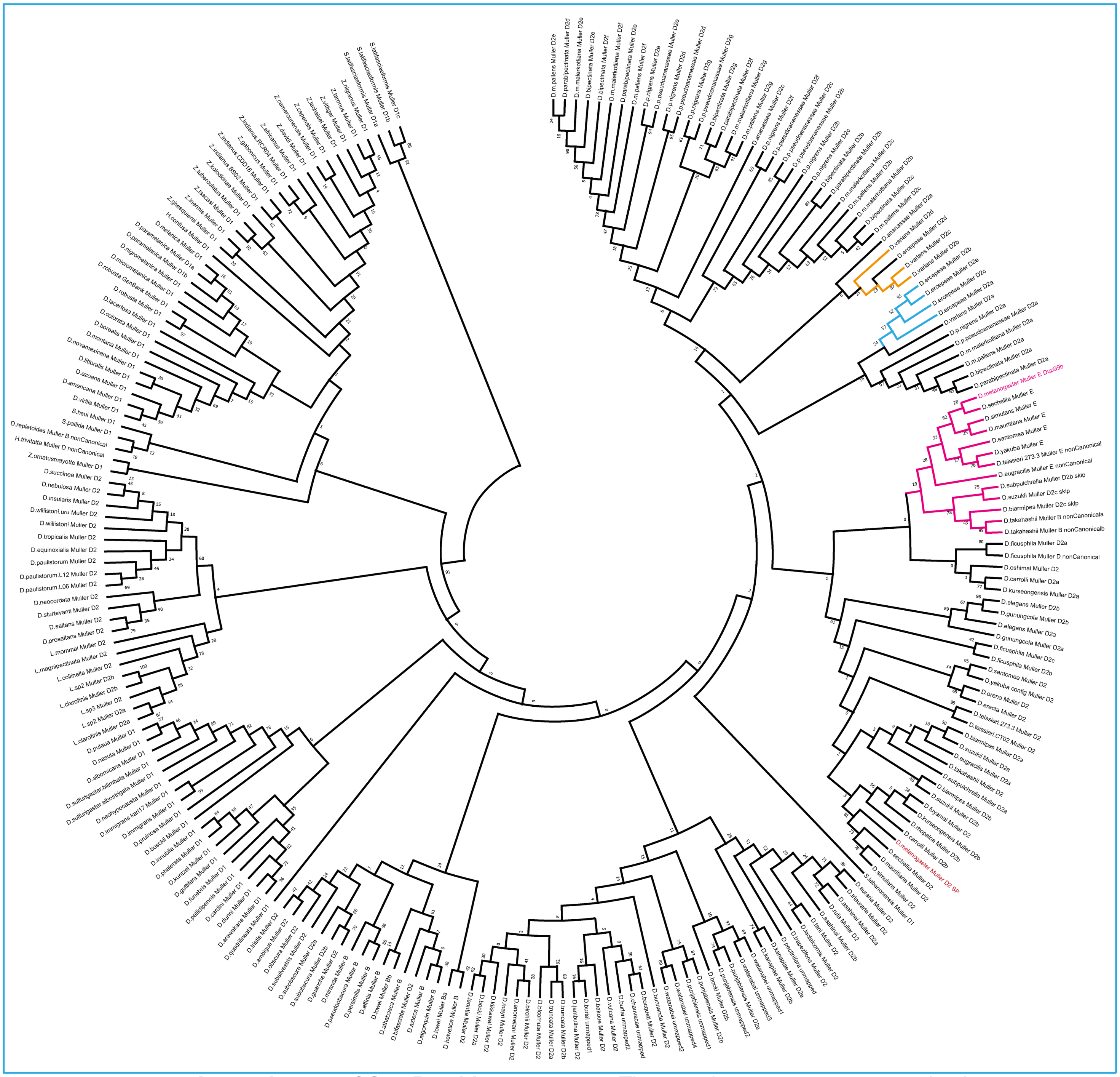

**Fig. S4. A protein tree of Sex Peptide sequences.** The protein tree was constructed using the Maximum Likelihood method on a MUSCLE alignment of 233 extracted SP amino acid sequences. The sequences include the signal peptide. Several clades of interest that are discussed in the text are coloured. A higher resolution figure can be downloaded at: https://osf.io/f7n6p.

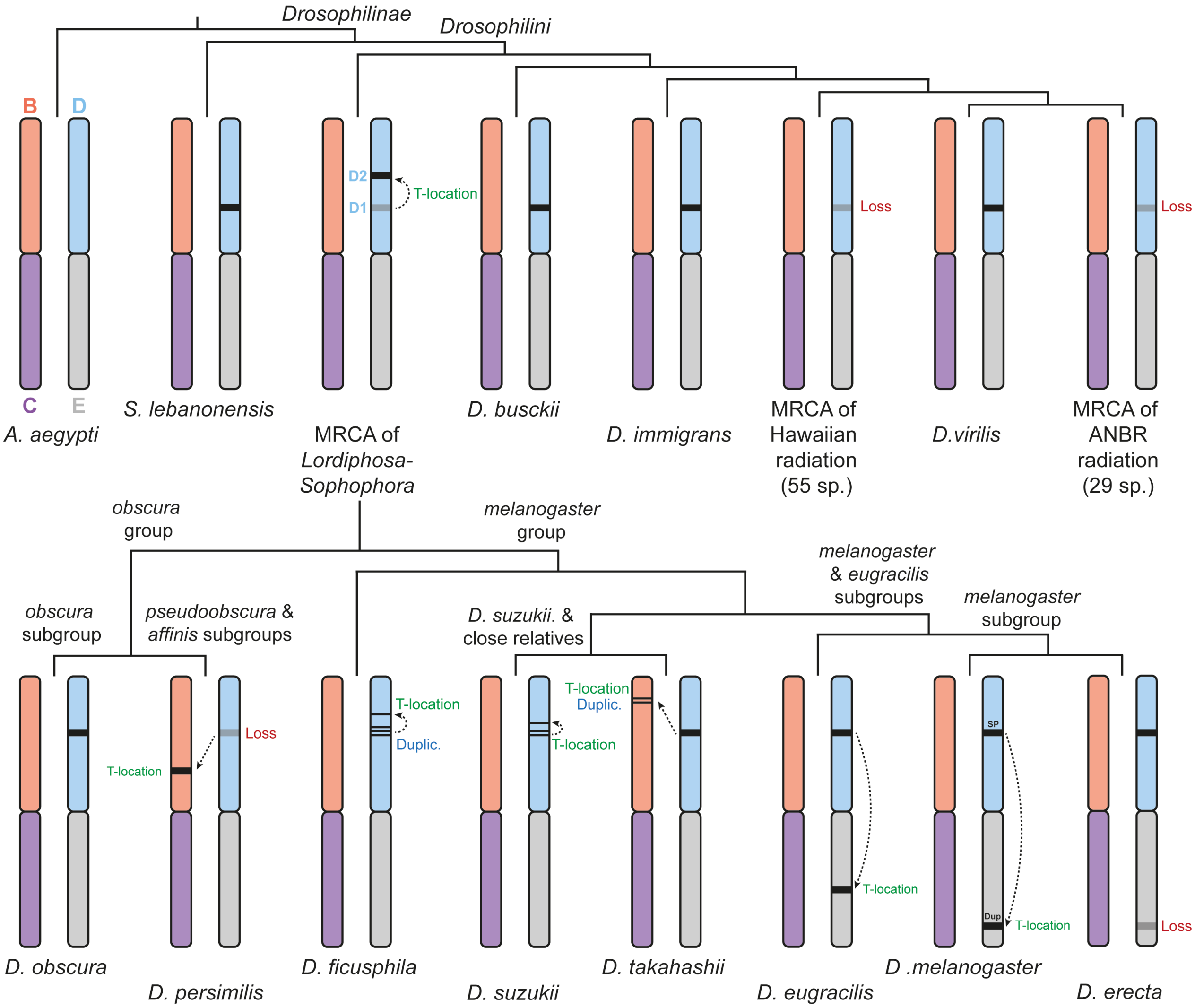


**Fig. S5. An overview of duplications, losses, and translocations of *Sex Peptide*.** This schematic summarises several translocations, duplications, and losses that have occurred within the *Drosophilinae*. Muller elements B to E are depicted for each species. Ancestrally, *SP* appears to be present within a gene neighbourhood ('Muller D1') that maps to Muller element D in *D. melanogaster* and contains orthologs of *FoxK, mRpL2,* and *NaPi-III.* In the most recent common ancestor (MRCA) of the *Sophophora-Lordiphosa* radiation, *SP* has been lost from Muller D1 and has translocated to a new position that maps to the same Muller element ('Muller D2'). This Muller D2 neighbourhood contains orthologs of *capricious, CG14111,* and *CG17687.* In the other *Drosophilini* lineage, *SP* is retained in the original genomic location, with the exceptions of *Hirtodrosophila trivittata* and *D. repletoides*, and several independent losses, which includes the Hawaiian *Drosophila* and a monophyletic radiation covering the *annulimana*, *bromeliae*, *nannoptera* and *repleta* ('ANBR') groups.

Within the *Sophophora-Lordiphosa* radiation, *SP* has repeatedly duplicated and/or translocated to new locations. In the *obscura* group, *SP* is retained in the Muller D2 position that's ancestral to the *Sophophora-Lordiphosa* radiation in all species except those within the *pseudoobscura* and *affinis* subgroups. In these subgroups, *SP* has been lost from Muller D2 and translocated to a neighbourhood that maps to *D. melanogaster* Muller element B and which contains orthologs of *pgant35A* and *spel1*.

Within the *melanogaster* group, SP has repeatedly and independently translocated and duplicated. Members of the *melanogaster* subgroup retain *SP* in Muller D2, but show a translocated copy (*Dup99b*) on Muller E. In several species, including *D. erecta* and *D. orena,* the Muller E copy has been lost. *D. eugracilis,* the outgroup to the *melanogaster* subgroup, shows a Muller D2 copy but its *Dup99b* copy maps to a non-canonical neighbourhood on Muller E. *D. suzukii* and its close relatives (*D. biarmipes* and *D. subpulchrella*) have 1 to 2 copies of *SP* in Muller D2 with an additional copy just the other side of the D2 neighbourhood; *D. takahashii* has a single copy in D2 with a translocation to and subsequent duplication in Muller B. A protein tree supports a *Dup99b* identity for the copies that fall outside of the canonical Muller D2 location in *D. eugracilis, D. suzukii, D.biarmipes, D. subpulchrella,* and *D. takahashii* (Fig. S4). *D. ficusphila* has 3 copies in D2 with an additional copy in a different neighbourhood on Muller element D that contains orthologs of *CG11905, CG13023,* and *CG13028*. Where multiple SP copies are present within a small area, we have thinned the bar signifying an SP gene. Positions of SP genes on each Muller element are approximate and not to scale. Not shown is a translocation of one of the *montium* subgroup species' *D. kanapiae* two copies to a novel gene neighbourhood on Muller D, nor a translocation to Muller E in *D. teissieri* (see Fig. S7 legend). This neighbourhood is ~300Kb from Muller D2 in *D. melanogaster.* A higher resolution figure can be downloaded at: https://osf.io/6ty2n.

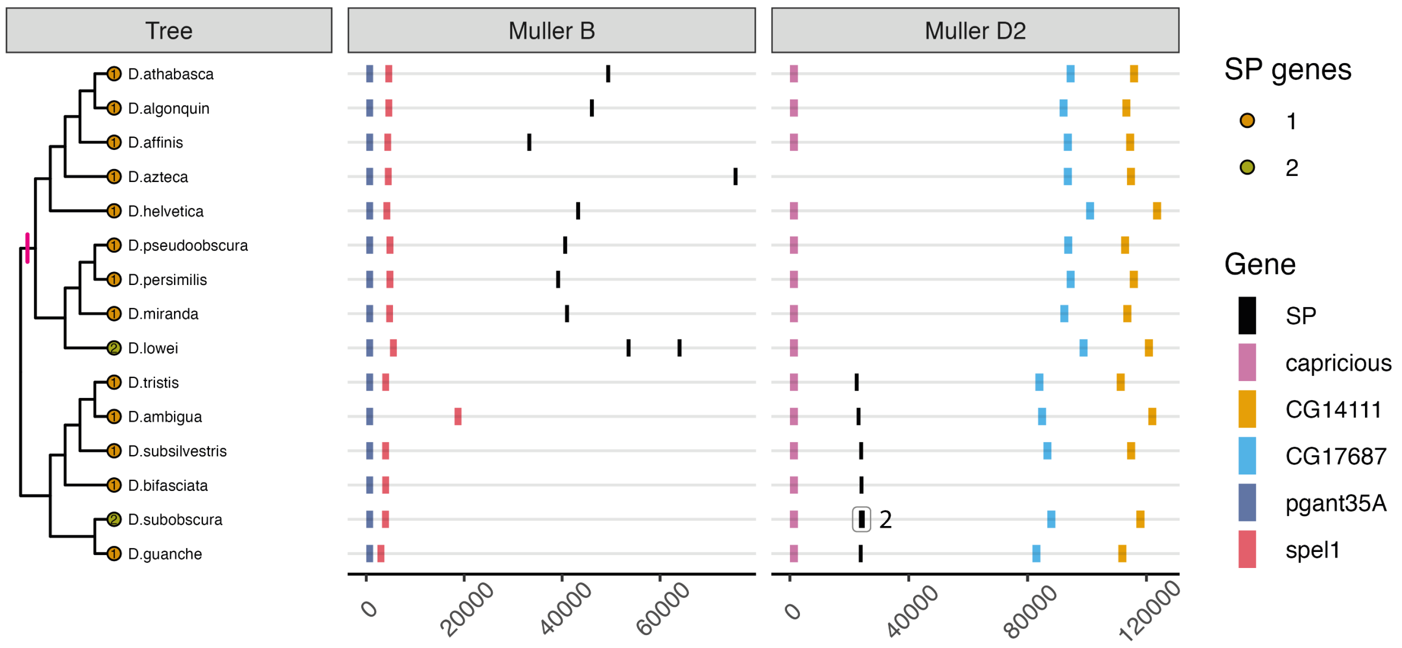


**Fig. S6.** ***Sex Peptide*** **translocated at the base of the clade that includes the** ***affinis*** **and** ***pseudoobscura*** **subgroups.** The number of *SP* genes identified in each species is given at the tip of each branch. *SP* genes are plotted in relation to one of two gene neighbourhoods, one of which mapped to Muller element B in *D. melanogaster* and the other of which mapped to Muller element D. This 'Muller D2' neighbourhood represents the canonical location of *SP* within the *Sophophora-Lordiphosa* radiation. Note that the absence of neighbourhood genes in a species doesn't necessarily indicate their absence from that species' genome. It may reflect the presence of a contig breakpoint within the neighbourhood that prevented us from resolving the neighbourhood's structure. Note also the apparently independent duplications of *SP* in *D. lowei* and *D. subobscura.* In the latter case, the two bars are hard to visually separate due to their close proximity. A pink line indicates the branch where the translocation occurred. A higher resolution figure can be downloaded at: https://osf.io/kn53f.

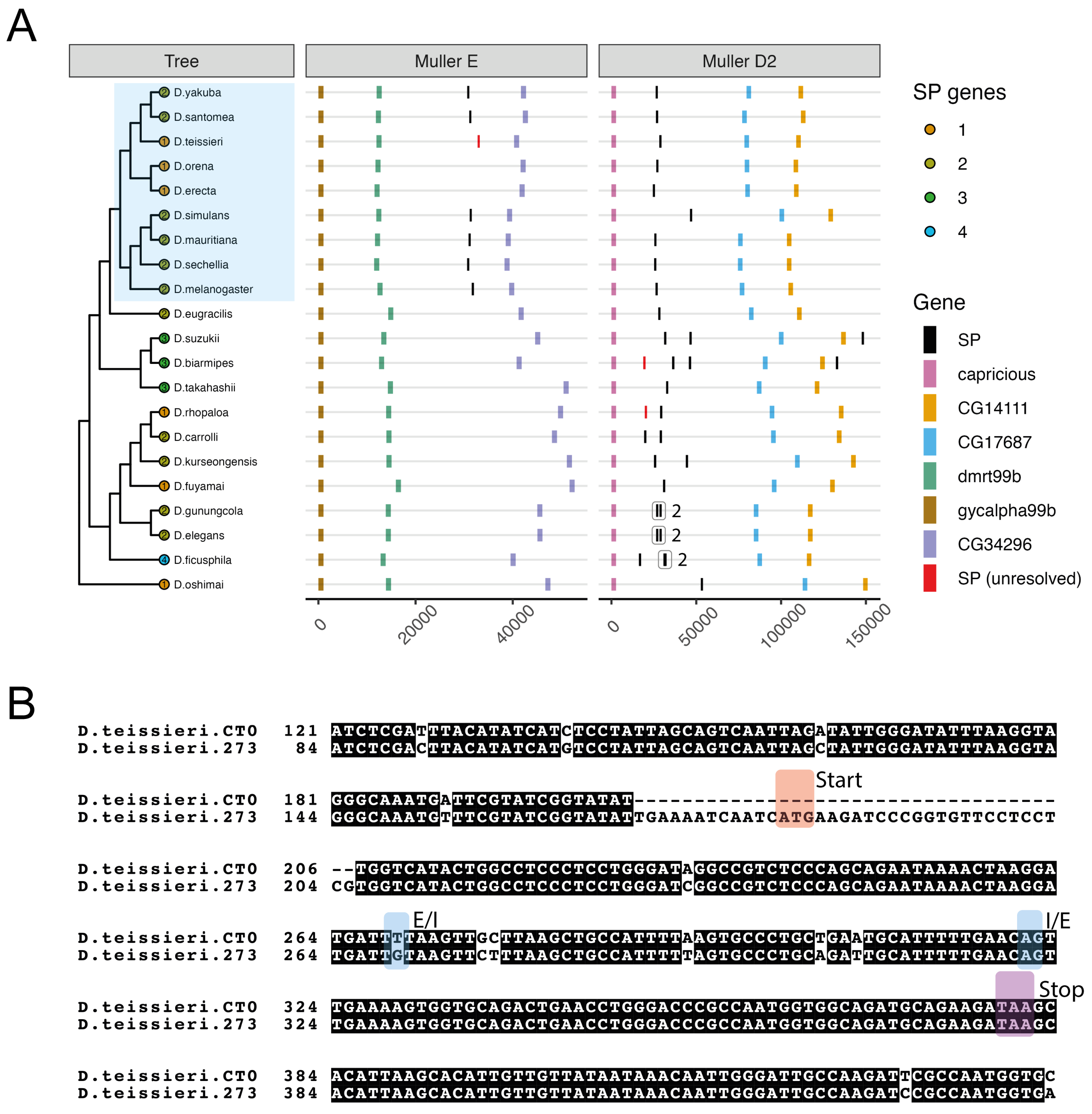


**Fig. S7. The localisation of a *Sex Peptide* gene to the *Dup99b* position is restricted to the *melanogaster* subgroup**. (A) A phylogeny of the "Oriental" lineage species used in this study. Species in the *melanogaster* subgroup are highlighted with a blue square. The number of *SP* genes identified in each species is given at the tip of each branch. 'Unresolved' SP sequences, shown in red, are those that passed the reciprocal blast test and fell within the syntenic Muller D2 gene neighbourhood but for which we could not resolve an SP-like protein structure (*e.g.,* due to a premature stop codon). The structures of the Muller E and Muller D2 gene neighbourhoods are plotted on the right-hand side of the figure. Several sequences are not plotted as they fall outside of these neighbourhoods: one from *D. ficusphila*, which falls in a separate neighbourhood that maps to Muller element D; one from *D. eugracilis,* which maps to a region on Muller element E near to, but distinct from, the canonical *Dup99b* position; and two from *D. takahashii,* which map to Muller element B. (B) Alongside *SP* genes in the canonical Muller D2 position, both strains of *D. teissieri* (CT02 and 273.3) also encoded an additional copy in a different gene neighbourhood that mapped to Muller element E in *D. melanogaster* and contained orthologs of *poly, Lip3,* and *mthl12*. This neighbourhood was ~16.3Mb away from *Dup99b* on *D. melanogaster* Muller element E. However, we could only resolve an SP-like protein sequence for this copy in one of the two strains (273.3). CT02 contained a 37bp deletion covering the start codon, as shown in this alignment. E/I and I/E represent the exon/intron boundaries. A higher resolution figure can be downloaded at: https://osf.io/kmy83.

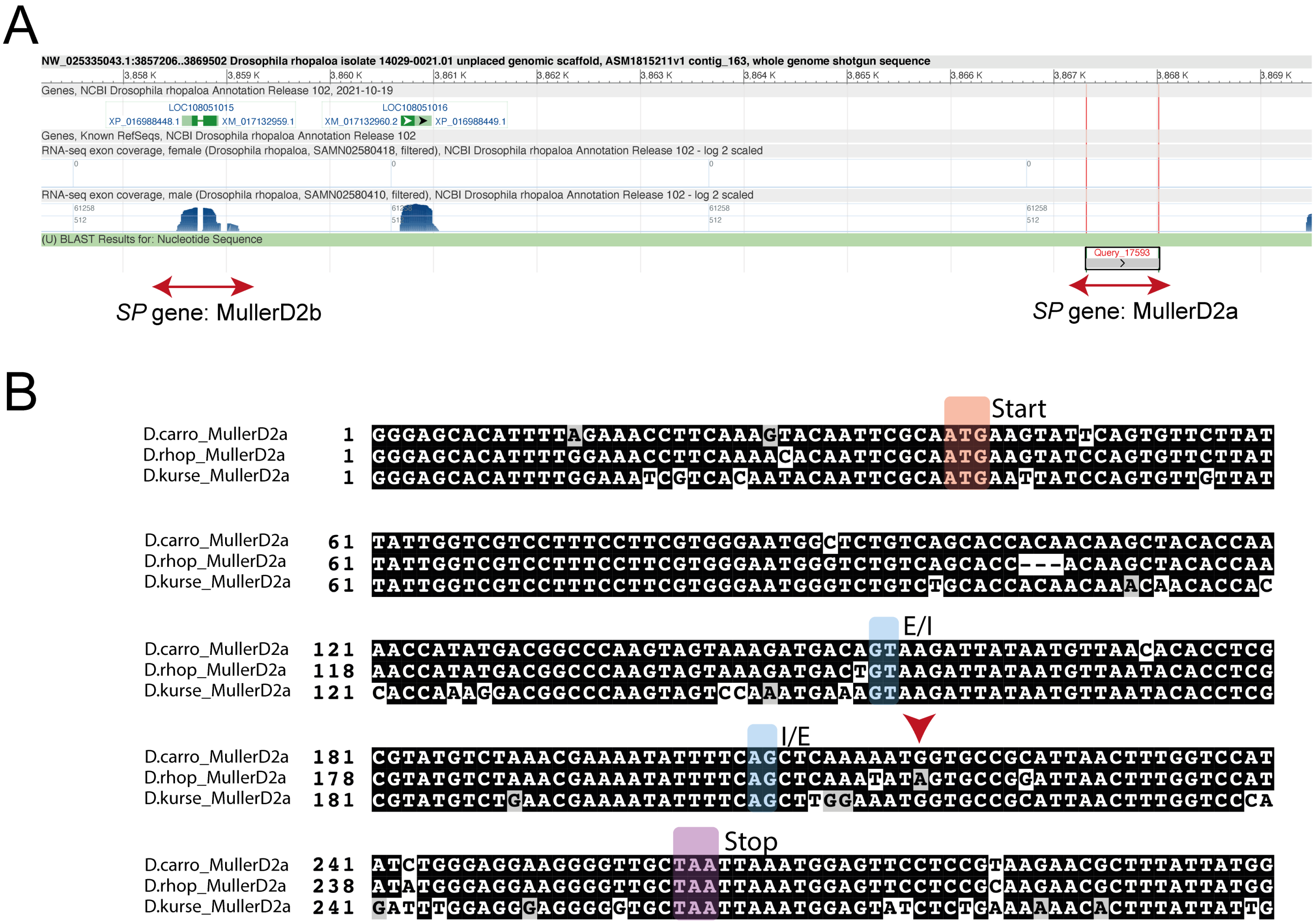


**Fig. S8. Pseudogenisation of a *Sex Peptide* paralog in *D. rhopaloa*.** As in its two closest relatives, *D. carrolli* and *D. kurseongensis,* we detected two *SP* copies in the Muller D2 neighbourhood of *D. rhopaloa.* (A) In this screenshot from the NCBI genome data browser we can see the position of the two *SP* genes (depicted by red arrows) in the genome we searched. One copy, which we refer to as 'MullerD2b', corresponds to the gene LOC108051015. RNA-seq tracks from male and female samples show clear male-specific expression spanning the two exons that correspond to this gene. The other *SP* gene we identify, the copy closest to *capricious* and which we refer to as 'MullerD2a', has neither evidence of expression nor an associated annotated feature. Interestingly, LOC108051016, which separates the two *SP* hits, showed male-specific expression and encoded a protein with no clear homologous sequence in the *D. melanogaster* reference protein list, suggesting it may encode a lineage- and sex-specific short peptide. (B) Along with the absence of expression, we couldn't resolve a SP-like protein sequence for *D. rhopaloa*_MullerD2a. This was due to a G to A point mutation that introduced a premature stop codon into the SP reading frame, the position of which is indicated by a red arrow. E/I and I/E represent the exon/intron boundaries. A higher resolution figure can be downloaded at: https://osf.io/yftgw.

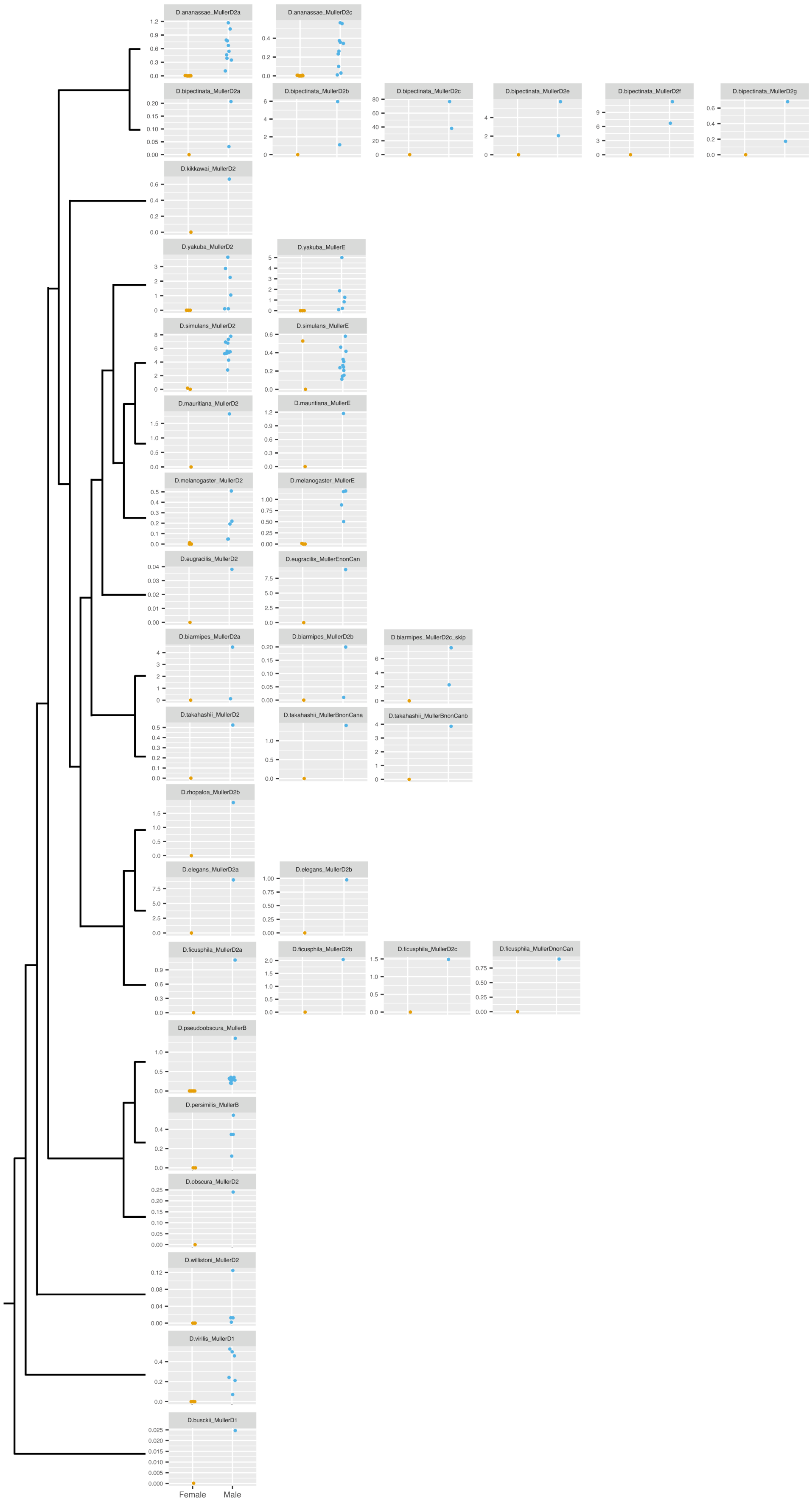
D.ananassae_MullerD2a
D.ananassae_MullerD2c
D.bipectinata_MullerD2a
D.bipectinata_MullerD2b
D.bipectinata_MullerD2c
D.bipectinata_MullerD2e
D.bipectinata_MullerD2f
D.bipectinata_MullerD2g
D.kikkawai_MullerD2
D.yakuba_MullerD2
D.yakuba_MullerE
D.simulans_MullerD2
D.simulans_MullerE
D.mauritiana_MullerD2
D.mauritiana_MullerE
D.melanogaster_MullerD2
D.melanogaster_MullerE
D.eugracilis_MullerD2
D.eugracilis_MullerEnonCan
D.biarmipes_MullerD2a
D.biarmipes_MullerD2b
D.biarmipes_MullerD2c_skip
D.takahashii_MullerD2
D.takahashii_MullerBnonCana
D.takahashii_MullerBnonCanb
D.rhopaloa_MullerD2b
D.elegans_MullerD2a
D.elegans_MullerD2b
D.ficusphila_MullerD2a
D.ficusphila_MullerD2b
D.ficusphila_MullerD2c
D.ficusphila_MullerDnonCan
D.pseudoobscura_MullerB
D.persimilis_MullerB
D.obscura_MullerD2
D.willistoni_MullerD2
D.virilis_MullerD1
D.busckii_MullerD1
Female
Male

**Fig. S9. Male-biased expression is a conserved feature of *SP* genes.** We obtained sex-labelled RNA-seq datasets from whole body samples of 19 different species hosted on NCBI. We extracted the maximum exonic read coverage value for each *SP* locus and divided it by the equivalent value extracted from the housekeeping gene *RpL32* to normalise expression across samples. Each plot depicts data from a different *SP* gene, with rows corresponding to copies within the same species. Male samples are plotted in blue and female samples in orange. Genes are named by species, the Muller element, and neighbourhood they map to, and, in the case of tandemly arranged copies in the Muller D2 neighbourhood, in their order from *capricious.* The one *SP* copy that does not show clear male-biased expression mapped to the Muller element E *Dup99b* location in *D. simulans.* However, this sample (SAMN02713493) was almost certainly contaminated with males or its sex mislabelled as, unlike the other female sample, it showed appreciable expression of the male-specific long non-coding RNAs *roX1* and *roX2*, which function in X chromosome dosage compensation (Fig. S10). A higher resolution figure can be downloaded at: https://osf.io/9mqyb.

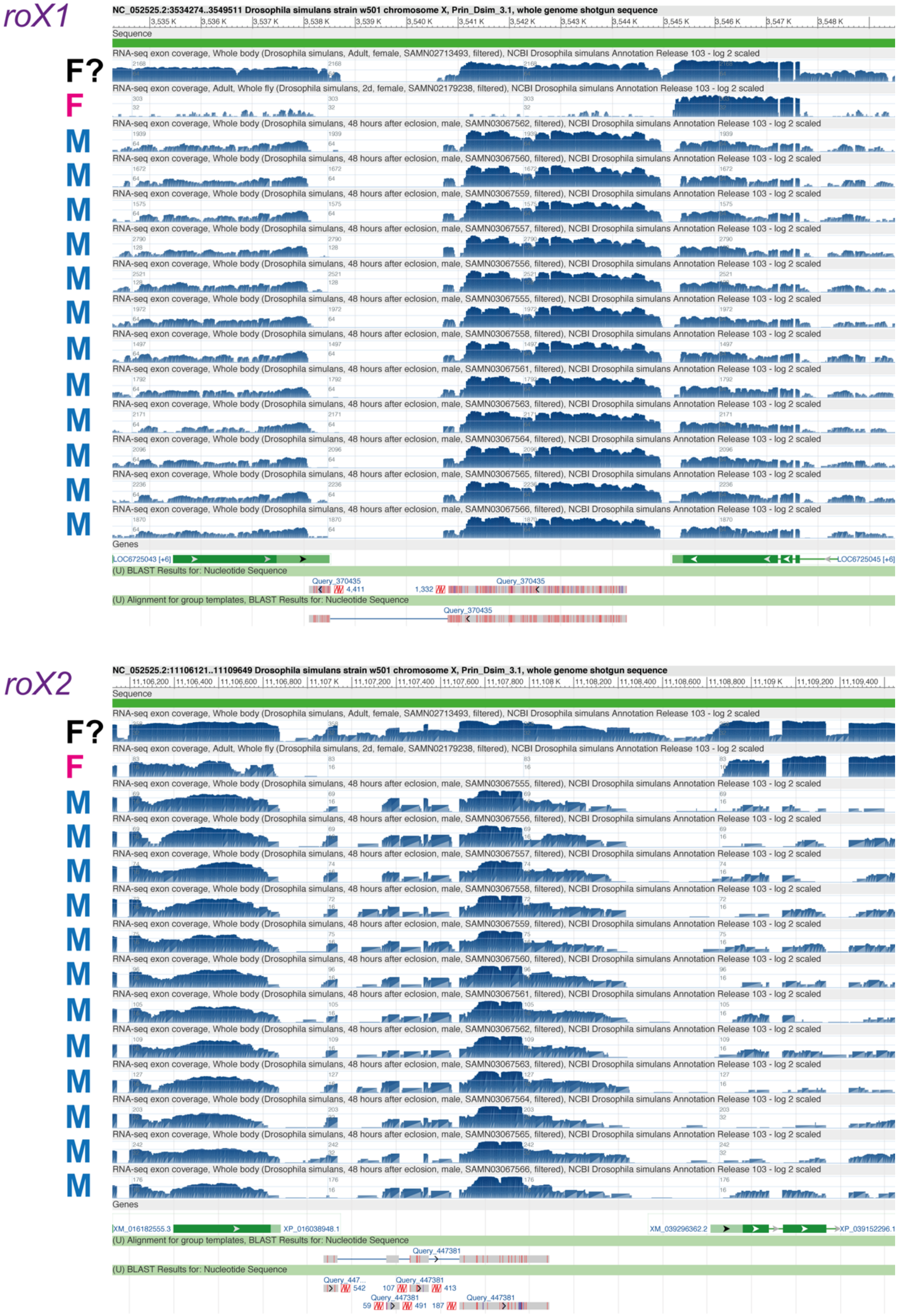


**Fig. S10. Expression of *SP* in a whole-body *D. simulans* female sample is likely due to contamination.** In Fig. S9, we showed that a single RNA-seq sample from a female of one of the

19 species examined showed appreciable *SP* expression. This sample, however, was almost certainly mislabeled or contaminated with males. To establish this, we used BLAST to query the *D. melanogaster* transcript sequences of *roX1* (FBgn0019661) and *roX2* (FBgn0019660) against the *D. simulans* reference genome (Prin_D.sim_3.1) and then overlaid over the returned hit the RNA-seq expression tracks from the whole-body male and female samples plotted for *D. simulans* in Fig. S10. For both genes, we detect clear, male-like expression in the *SP*-expressing female sample SAMN02713493, but not the other female sample. Among adults, *roX1* and *roX2* are exclusively expressed in males (70–72). M = male, F = female. F? = the female-labeled SAMN02713493 sample. A higher resolution figure can be downloaded at: https://osf.io/8zscm.

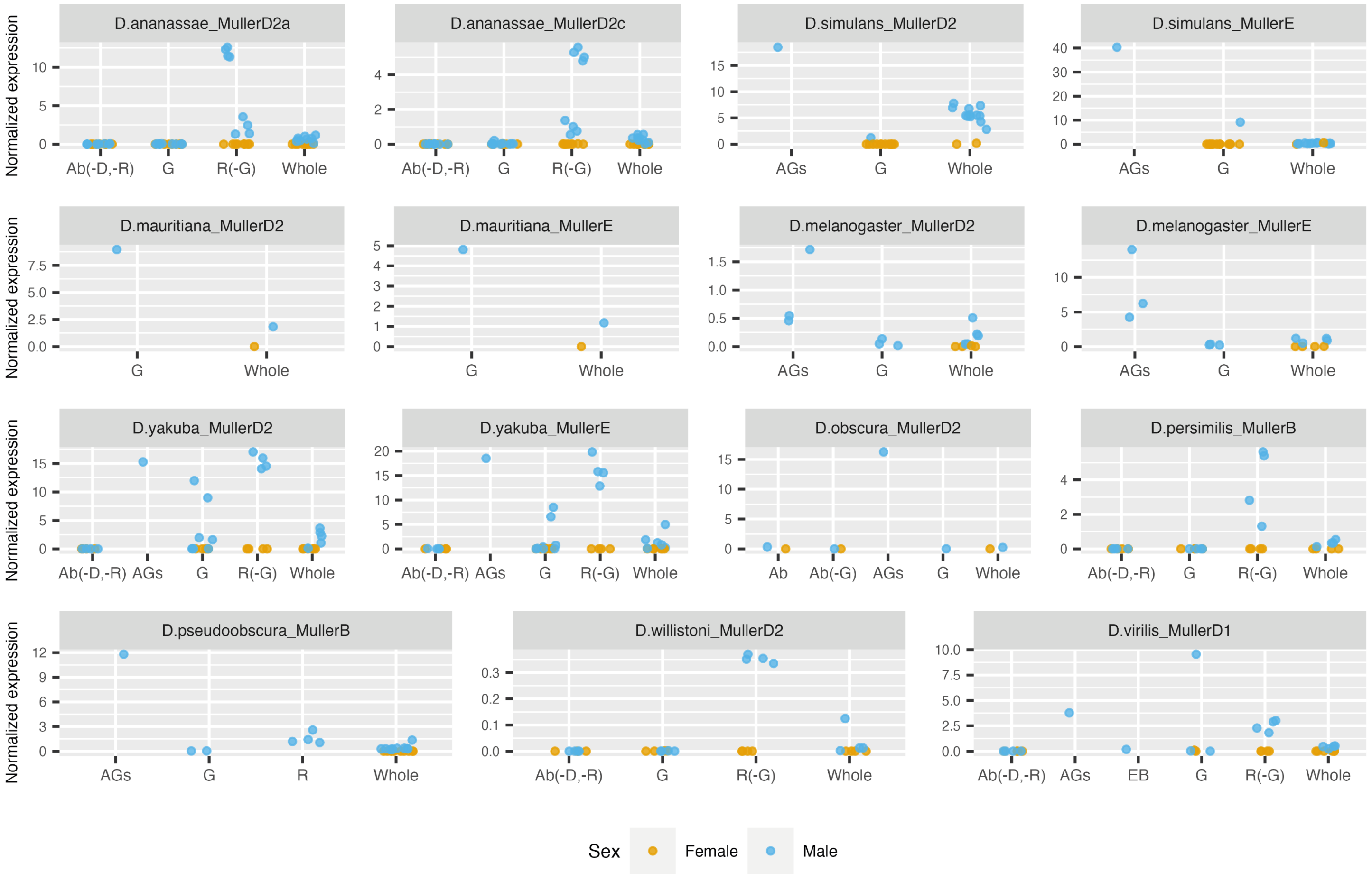


**Fig. S11. Male reproductive tract biased expression is a conserved feature of *SP* genes.** We obtained tissue-specific RNA-seq datasets from 10 different species hosted on NCBI. We extracted the maximum exonic read coverage value for each *SP* locus and divided it by the equivalent value extracted from the housekeeping gene *RpL32* to normalise expression across samples. Each plot depicts data from a different *SP* homolog. Genes are named by species, the Muller element, and neighbourhood they map to, and, in the case of tandemly arranged copies in the Muller D2 neighbourhood, in their order from *capricious.* Note that there is no D.ananassae_MullerD2b as this is the 'unresolved' copy that can be seen in Figure 4, for which we failed to find expression in any examined tissue. Ab (-D, -R) = Abdomen, digestive and reproductive tracts removed; R = Reproductive tract; G = Gonad (ovaries/testes); R (-G) = Reproductive tract without gonad; Whole = Whole flies; AGs = Accessory glands; Ab = Abdomen; Ab (-G) = Abdomen without gonad; EB = Ejaculatory bulb. The raw data, including dataset accessions, is available in https://osf.io/ydpfz. A higher resolution figure can be downloaded at: https://osf.io/zguer.

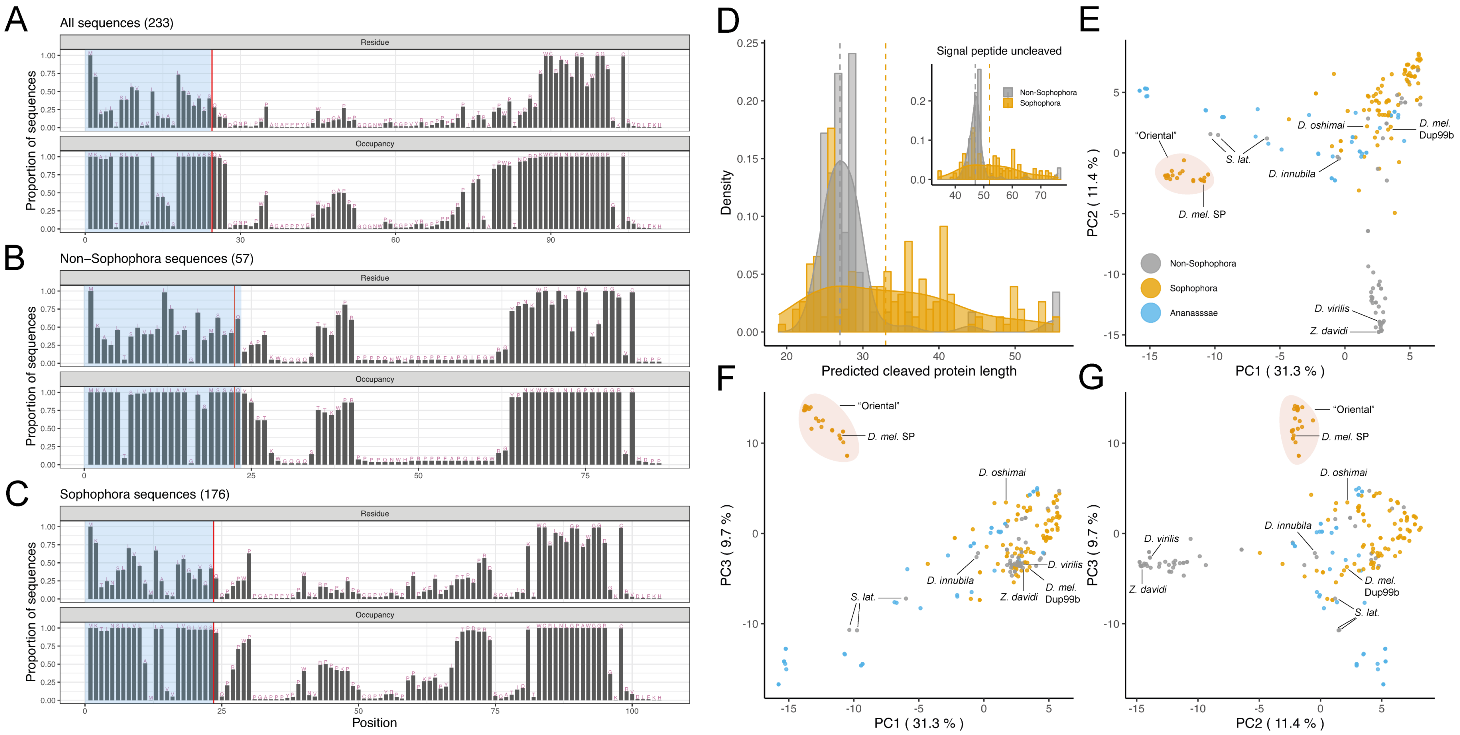


**Fig. S12. Sex Peptide has undergone domain-specific evolutionary change.** (A-C) Consensus sequences based on MAFFT alignment of the resolvable amino acid sequences of (A) 233 SP genes, (B) 57 from the non-*Sophophora-Lordiphosa* species, and (C) 176 from the *Sophophora-Lordiphosa* species. In each panel, the top plot gives the proportion of sequences with the consensus amino acid in the same position, while the bottom plot gives the proportion of sequences in which each position is occupied in the alignment. A red line indicates the position of a predicted signal peptide cleavage site determined using SignalP 6.0. The probability of a cleavage site in the given position was 0.980 in A, 0.973 in B, and 0.982 in C. (D) The distribution of predicted protein lengths for each resolved SP sequence plotted separately for *Sophophora-Lordiphosa* and non-*Sophophora-Lordiphosa* sequences. The main panel shows the protein lengths after removing the signal peptide at the predicted site; the inset shows the full lengths prior to cleavage of the predicted peptide. Four sequences, two from each of the *montium* subgroup species *D. punjabiensis* and *D. watanabei,* did not contain a predicted signal peptide and are therefore omitted from the post-cleavage plot. Dashed lines indicate median lengths for the *Sophophora-Lordiphosa* and non-*Sophophora-Lordiphosa* species, separately. The smoothed curve shows a kernel density estimate, a smoothed version of the histogram, derived using the geom_density function in ggplot2. (E-G) PCA plots based on BLOSUM62 substitution scores from the MAFFT-aligned resolved SP sequences. (E) PC1 vs PC2. (F) PC1 vs PC3. (G) PC2 vs PC3. The percentage values in the axis titles reflect the proportion of variance explained by a given PC. Points are coloured based on whether they correspond to *Sophophora-Lordiphosa,* non-*Sophophora-Lordiphosa,* or *ananassae* subgroup species. A higher resolution figure can be downloaded at: https://osf.io/js97v.

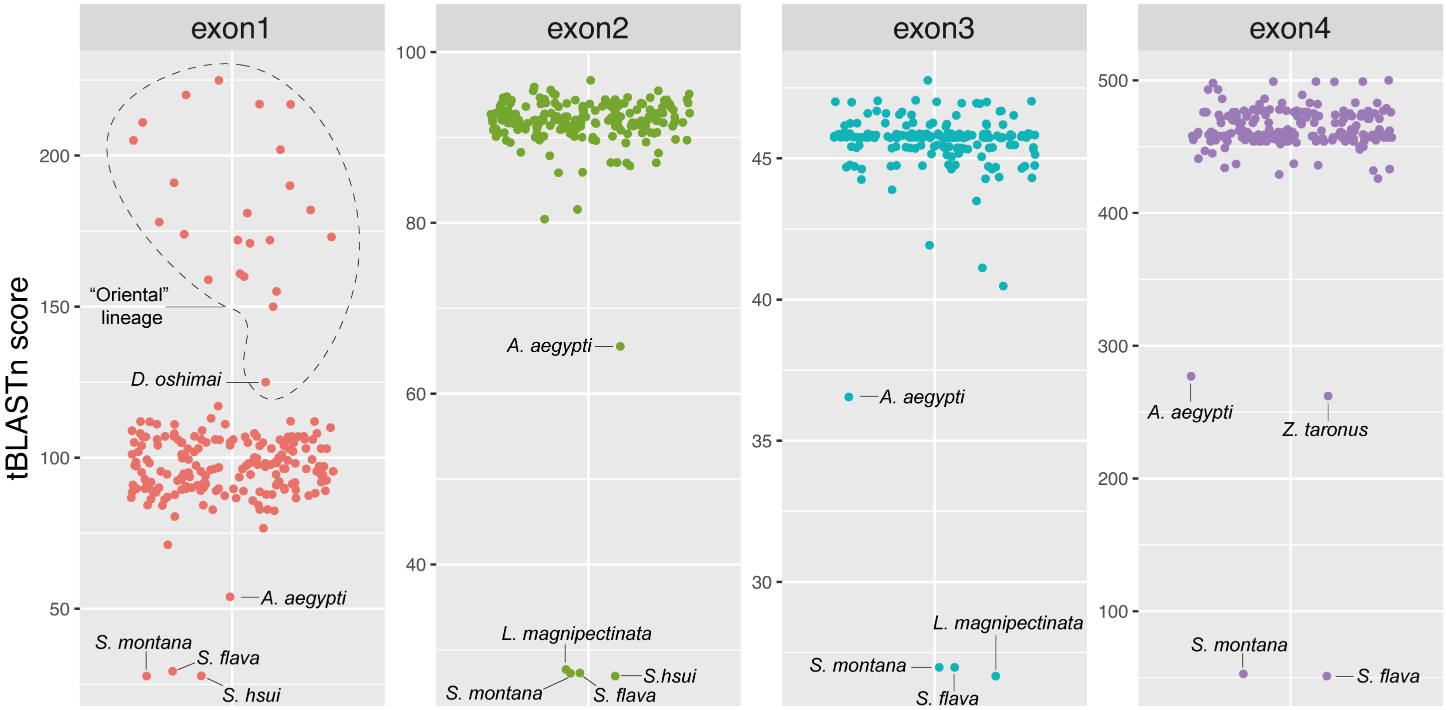


**Fig. S13. The protein sequence encoded by exon 1 shows the greatest between-species variation among the exons of *SPR*.** The tBLASTn scores plotted from the top hit returned when the amino acid sequence encoded by each of *D. melanogaster's* four *SPR* exons was queried against each species in our dataset. *A. aegypti* is the most phylogenetically distant species in the dataset and its position is labelled in each plot. The circled region in the exon 1 plot contains all species in our dataset from the “Oriental” lineage. Note the sharp, exon-specific drop in score that occurs outside of this lineage. All other labelled points correspond to the 5 species in which we couldn't resolve SPR sequences (*Scaptomyza montana, S. flava, S. hsui, Lordiphosa magnipectinata,* and *Zaprionus taronus*). Note how some species have low scores recorded for each exon (*e.g. S. montana* and *S. flava*), indicative of loss of the whole gene, others show a drop in a subset of exons, such as *Z. taronus*, which recorded a drop in score only in exon 4 (due to a premature stop codon). A higher resolution figure can be downloaded at: https://osf.io/wr4ds.

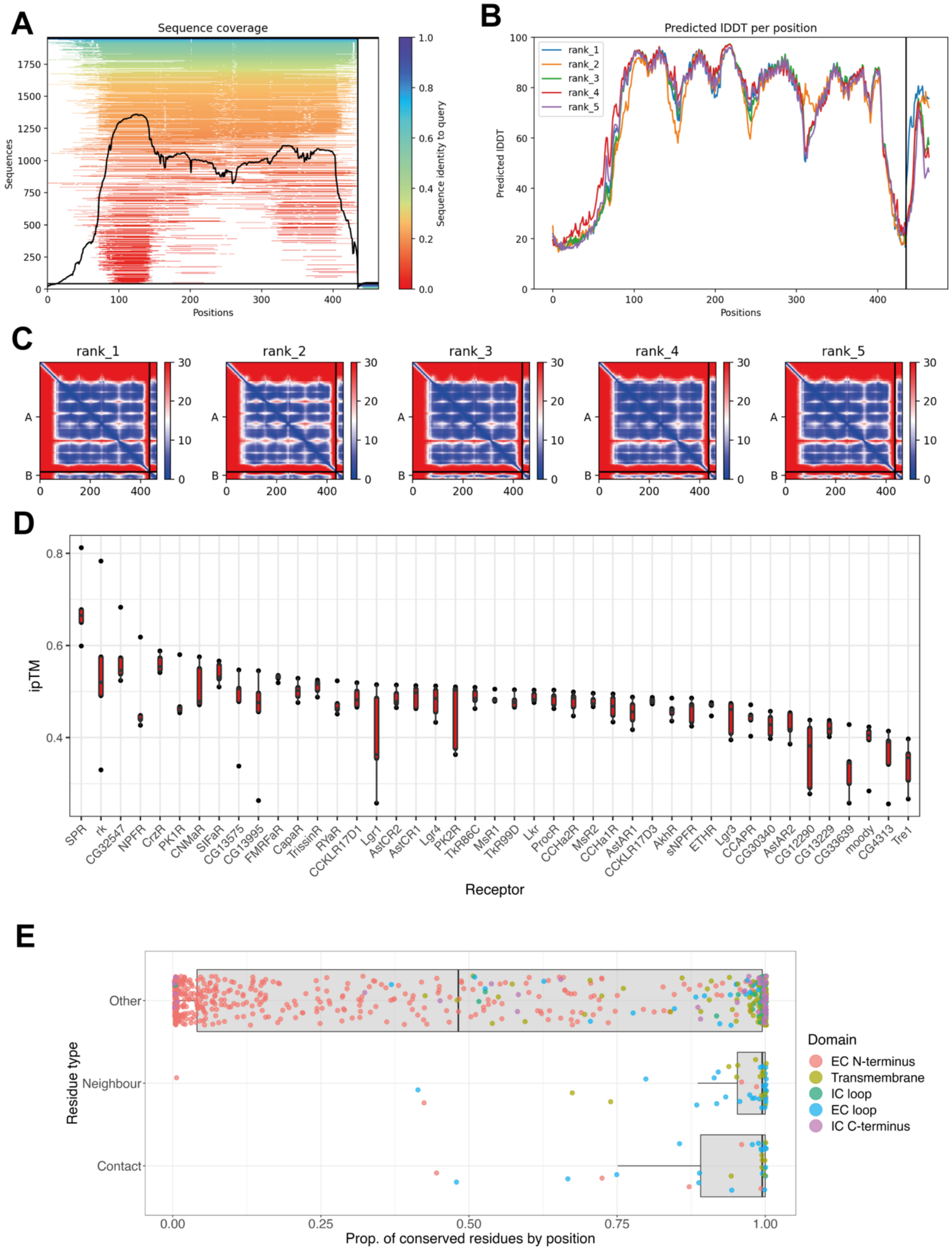


**Fig. S14. Summary statistics for the ColabFold-predicted interactions between SPR and SP.** (A-C) Summary plot outputs from the ColabFold prediction of SP-SPR interactions. (A) The multiple sequence alignment coverage per position. The vertical black line on the right-hand side of the image marks the separation between SP and SPR. (B) The AlphaFold confidence measure lDDT (local Distance Difference Test), which provides a measure of the model confidence at each position. (C) The predicted alignment error per position for SPR ('A') and SP ('B') for each of the 5 ranked structures (rank 1 is the highest, *i.e.,* best performing). (D) The ipTM scores, which describes the confidence in the interface structure, plotted for each of the 5 ranked structures predicted between SP and each *D. melanogaster* Class A G-protein coupled receptor. (E) A boxplot showing the proportion of sequences at each position in an alignment of 193 N-terminus trimmed SPR sequences that share the consensus residue. Residues are plotted separately based on whether they are a predicted SP-contacting residue, a residue that flanks (either side) a predicted contact residue, or neither. Points are coloured based on the SPR domain they belong to. EC = extracellular; IC = intracellular. A higher resolution figure can be downloaded at: https://osf.io/3hq47.

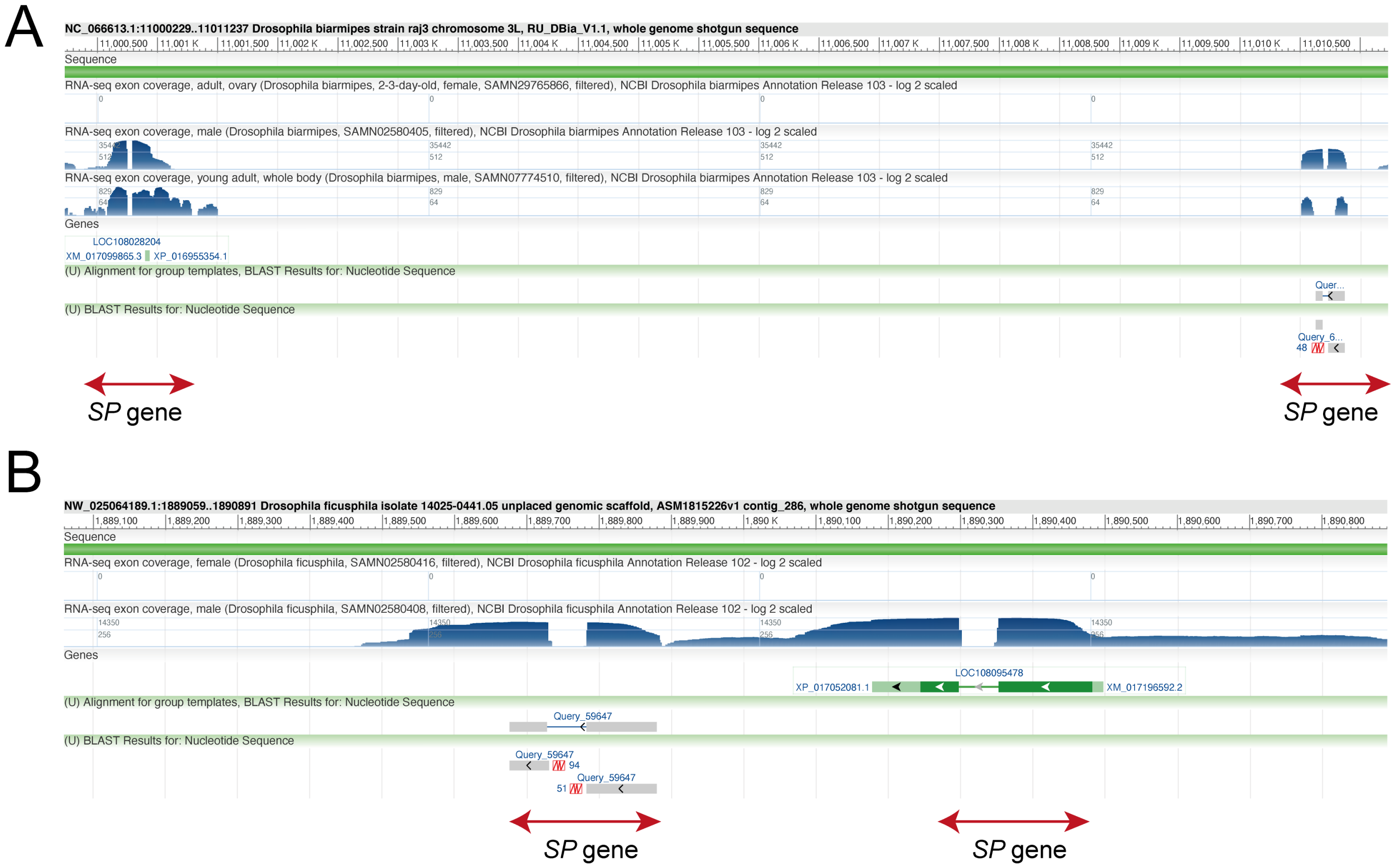


**Fig. S15. Unannotated *Sex Peptide* genes in *D. biarmipes* and *D. ficusphila*.** Screenshot from the NCBI genome data browser showing the positions of *SP* hits (red arrows) in genomes of *D. biarmipes* (A) and *D. ficusphila* (B). In each case, the *SP* genes we identify are accompanied by male-specific expression in the RNA-seq exon coverage tracks, but only one of each species' two shown hits has an accompanying feature annotation in the 'Genes' window. The 'BLAST' window shows the position of the unannotated SP gene in each species. A higher resolution figure can be downloaded at: https://osf.io/tyxqn.

**Movie S1 (separate file).** SPR coloured by sequence conservation at each residue (high values = high conservation).

**Movie S2 (separate file).** The predicted interface between SPR (beige) and SP (green). Pink residues in SPR indicate predicted contact residues. Red dotted lines indicate predicted H-bond-forming residues between SP and SPR.

**Movie S3 (separate file).** The predicted interface between SPR and SP. Interfacing residues are coloured.

**Movie S4 (separate file).** The predicted interface between SPR (beige) and SP (green). Pink residues in SPR indicate the 10 residues identified using MEME as showing evidence of episodic positive selection.
